# A Brownian ratchet model for DNA loop extrusion by the cohesin complex

**DOI:** 10.1101/2021.02.14.431132

**Authors:** Torahiko L Higashi, Minzhe Tang, Georgii Pobegalov, Frank Uhlmann, Maxim Molodtsov

## Abstract

The cohesin complex topologically encircles DNA to promote sister chromatid cohesion. Alternatively cohesin extrudes DNA loops, thought to reflect chromatin domain formation. Here, we propose a structure-based model explaining both activities, supported by biochemical experiments. ATP and DNA binding to cohesin promote conformational changes that guide DNA through a kleisin gate into a DNA gripping state. Two HEAT-repeat DNA binding modules, associated with cohesin’s heads and hinge, are now juxtaposed. ATP hydrolysis disassembles the gripping state, allowing unidirectional hinge module movement to complete topological DNA entry. Without initial kleisin gate passage, biased hinge module motion during gripping state resolution creates a Brownian ratchet that drives loop extrusion. Molecular-mechanical simulations of gripping state formation and resolution cycles recapitulate experimentally observed DNA loop extrusion characteristics. Our model extends to asymmetric and symmetric loop extrusion, as well as z-loop formation. Loop extrusion by biased Brownian fluctuations has important implications for chromosomal cohesin function.

## Introduction

Cohesin is a member of the Structural Maintenance of Chromosomes (SMC) family of ring-shaped chromosomal protein complexes that are central to higher order chromosome organization (*Uhlmann, 2016*). Cohesin holds together replicated sister chromatids from the time of their synthesis in S phase until mitosis to ensure their faithful segregation during cell divisions (*Guacci et al., 1997; Michaelis et al., 1997*). In addition, budding yeast cohesin participates in mitotic chromosome condensation, while higher eukaryotic cohesin impacts gene regulation by defining boundary elements during interphase chromatin domain formation (*Parelho et al., 2008; Wendt et al., 2008*). Cohesin is also recruited to sites of double stranded DNA breaks to promote DNA repair by homologous recombination and to stalled DNA replication forks to aid restart of DNA synthesis (*Ström et al., 2004; Ünal et al., 2004; Tittel-Elmer et al., 2012*). Understanding how cohesin carries out all these biological functions requires the elucidation of the molecular mechanisms by which cohesin interacts with DNA, as well as how cohesin establishes interactions between more than one DNA.

Cohesin’s DNA binding activity is contained within its unique ring architecture (*Gligoris et al., 2014; Huis in ‘t Veld et al., 2014*). Two SMC subunits, Smc1^Psm1^ and Smc3^Psm3^, form long flexible coiled coils that are connected at one end by a dimerization interface known as the hinge (generic gene names are accompanied by fission yeast subunit names in superscript; fission yeast cohesin was used for the experiments and most of the structural analyses in this study). At the other end lie ABC transporter-type ATPase head domains whose dimerization is regulated by ATP binding. The two SMC heads are further connected by a kleisin subunit, Scc1^Rad21^, that completes the topological assembly of the cohesin ring. Two HEAT repeat subunits associate with the kleisin that promote topological loading of cohesin onto DNA. Scc3^Psc3^ binds to the kleisin middle region and forms a stable part of the complex. Scc2^Mis4^, together with its binding partner Scc4^Ssl3^, transiently associate with the kleisin upstream of Scc3^Psc3^. Once cohesin loading onto DNA is complete, Scc2^Mis4^-Scc4^Ssl3^ is replaced by a related HEAT repeat subunit, Pds5 (*Murayama and Uhlmann, 2015; Petela et al., 2018*). Because of its transient role, Scc2^Mis4^-Scc4^Ssl3^ is often thought of as a cohesin cofactor, termed ‘cohesin loader’ (*Ciosk et al., 2000; Murayama and Uhlmann, 2014*). Following topological loading, cohesin is free to linearly diffuse along DNA *in vitro* (*Davidson et al., 2016; Stigler et al., 2016*), while *in vivo* RNA polymerases push cohesin along chromosomes towards sites of transcriptional termination (*Lengronne et al., 2004; Davidson et al., 2016; Ocampo-Hafalla et al., 2016; Busslinger et al., 2017*). Cohesin promotes sister chromatid cohesion following DNA replication by topologically entrapping two sister DNAs (*Haering et al., 2008; Murayama et al., 2018*).

In addition to topologically entrapping DNA, *in vitro* experiments have revealed the ability of human cohesin to translocate along naked DNA in a directed motion, as well as its ability to extrude DNA loops (*Davidson et al., 2019; Kim et al., 2019*). These activities are reminiscent of those previously observed with a related SMC complex, condensin, a central mediator of mitotic chromosome formation (*Terakawa et al., 2017; Ganji et al., 2018*). Like topological loading onto DNA, loop extrusion by cohesin depends on its ATPase, as well as on the presence of the human Scc3^Psc3^ homolog SA1 and the cohesin loader (NIPBL-MAU2). In stark contrast to topological loading, cohesin is able to extrude DNA loops even if all three cohesin ring interfaces are covalently closed (*Davidson et al., 2019*). This suggests that loop extrusion does not involve topological DNA entry into the cohesin ring.

Several models have been proposed as to how SMC complexes extrude DNA loops. These include a tethered-inchworm model in which a scissoring motion of the ATPase heads translates into movement along DNA (*Nichols and Corces, 2018*). The DNA-segment-capture model instead suggests that transitions between open and closed configurations of the SMC coiled coils produce a pumping motion that constrains DNA loops (*Marko et al., 2019*). Finally, a scrunching model proposes that the SMC hinge reaches out to capture and reel in DNA loops (*Ryu et al., 2020b*). A characteristic of experimentally observed loop extrusion is that very small counterforces (< 1 pN) stall loop growth (*Ganji et al., 2018; Golfier et al., 2020*). Both the DNA-segment-capture and the scrunching model therefore foresee that diffusive DNA motion contributes to loop growth. In order to capture enlarging loops, both models assume complex coordination between DNA binding elements at the SMC heads and hinge, for which no experimental evidence as yet exists. Molecular details of the loop extrusion mechanism therefore remain elusive. Another important open question is whether SMC complexes can move along and extrude physiological chromatin substrates, decorated by histones and other DNA binding proteins.

We recently solved a cryo-EM structure of fission yeast cohesin with its loader in a nucleotide-bound DNA gripping state (*Higashi et al., 2020*). Together with DNA-protein crosslinking and biophysical experiments, this allowed us to trace the DNA trajectory into the cohesin ring by sequential passage through a kleisin N-gate and an ATPase head gate. We noticed, however, that kleisin N-gate passage might not be a strict prerequisite for DNA to reach the gripping state. We speculated that an alternative gripping state arises in which DNA has not passed the kleisin N-gate. While topological DNA entry is barred, this state might constitute an intermediate during loop extrusion. Here, we show how two DNA binding modules of the cohesin complex, formed by its HEAT repeat subunits (*Murayama and Uhlmann, 2014, 2015; Li et al., 2018; Collier et al., 2020; Kurokawa and Murayama, 2020*), are juxtaposed in the gripping state but swing apart following ATP hydrolysis. We illustrate how this swinging motion promotes completion of topological DNA entry or the beginning of loop extrusion, dependent on whether or not DNA passed the kleisin N-gate. Computational simulations of cycles of gripping state formation and resolution, in the latter scenario, demonstrate how a Brownian ratchet forms that can drive loop extrusion. Our analyses provide a molecular proposal for both topological entry into the cohesin ring as well as for DNA loop extrusion.

## Results

### Two DNA binding modules in the cohesin-DNA gripping state

Two cryo-EM structures of fission yeast and human cohesin with their cohesin loaders in a nucleotide-bound DNA gripping state revealed the presence of two DNA binding modules (*Higashi et al., 2020; Shi et al., 2020*). One is the DNA gripping interaction that forms as the cohesin loader, Scc2^Mis4^, clamps the DNA onto the engaged SMC ATPase heads (***Figure 1A***, left). Both Scc2^Mis4^ and the SMC heads contribute numerous positively charged surface residues that line the DNA path. Scc2^Mis4^ has undergone a marked conformational change compared to its crystallographically observed ‘extended’ conformation. Its N-terminal handle cradles the DNA in the gripping state, thereby adopting a ‘bent’ conformation. We refer to this composite DNA interaction site, consisting of Scc2^Mis4^ and the SMC ATPase heads, as the ‘Scc2-head module’.

**Figure 1.**
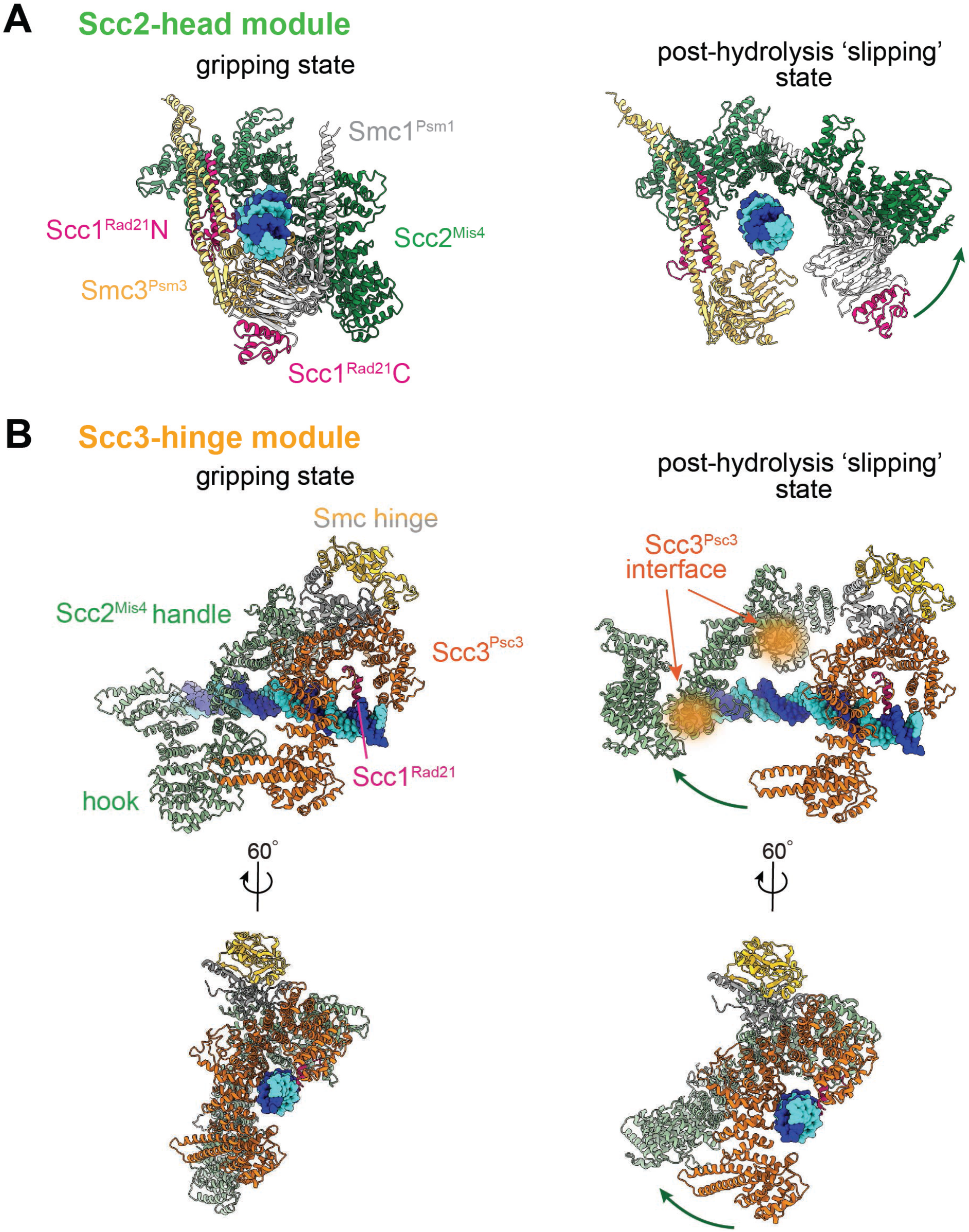
Two DNA binding modules in the gripping state and in their projected post-hydrolysis states. **(A)** The Scc2-head module in the gripping state and its predicted conformation following ATP hydrolysis. ATPase head disengagement and return of Scc2^Mis4^ from the gripping state conformation to its extended crystal structure form results in loss of DNA interactions, turning the module into its ‘slipping state’. **(B)** The Scc3-hinge module. The DNA binding surface of this module remains unaltered by ATP hydrolysis. However, the Scc2^Mis4^ conformational change following ATP hydrolysis leads to uncoupling of the Scc3-hinge from the Scc2-head module.

The second DNA contact is made by Scc3^Psc3^ in conjunction with the kleisin middle region (***Figure 1B***, left). EM and protein crosslink mass spectrometry analyses of the fission yeast gripping state place this module behind the cohesin loader. The SMC hinge touches down and bridges Scc3^Psc3^ and the cohesin loader, enabled by SMC coiled coil inflection at their elbows (***Figure 1 – figure supplement 1A***). The human gripping state cryo-EM structure shows Scc3^SA1^ in a similar orientation. The higher resolution of Scc3^SA1^ in this structure reveals details of its interaction with both the cohesin loader as well as the SMC hinge. For our further considerations of the gripping state, we model Scc3^Psc3^ to take the position of human Scc3^SA1^. The human structure shows Scc3^SA1^ and the kleisin middle region engaged with DNA, in a fashion similar to that previously seen in a crystal structure of budding yeast Scc3 bound to DNA (*Li et al., 2018*). The DNA interaction is again provided by an array of positively charged amino acids that line the combined Scc3^Psc3^ and kleisin surface. DNA-protein crosslink mass spectrometry data of the fission yeast gripping state further support this assignment (*Higashi et al., 2020*). We refer to this DNA binding site as the ‘Scc3-hinge module’.

### Predicted conformational changes following ATP hydrolysis

We now consider the consequences of ATP hydrolysis on the two DNA binding modules outlined above. Upon ATP hydrolysis, the composite DNA binding surface between the ATPase heads is disrupted, leading to loss of at least some of the DNA contacts within the Scc2-head module. If we further assume that Scc2^Mis4^ returns to its extended crystal structure form (*Higashi et al., 2020*), further DNA contacts are lost as the gripping state opens up (***Figure 1A***, right). We can see how, in this state, DNA is free to leave the Scc2-head module during topological DNA entry. The alternative kleisin path during loop extrusion (discussed below), prevents DNA from exiting the module but allows free DNA sliding movements in and out of the image plane. Because of the implications for loop extrusion, we refer to this shape of the Scc2-head module as its ‘slipping state’.

A bent Scc2^Mis4^ conformation that embraces DNA, compared to its extended crystal structure form, is common amongst the fission yeast, human and budding yeast gripping states (***Figure 1 – figure supplement 1B***)(*Collier et al., 2020; Shi et al., 2020*). This communality opens up the possibility that the gripping to slipping state conformational transition is a conserved feature of the Scc2-head module.

In contrast to the Scc2-head module, the DNA binding site in the Scc3^Psc3^-hinge module does not undergo an obvious conformational change when comparing its gripping and free crystal structure forms. This can be seen from the almost perfect overlap of human Scc3^SA1^ in the gripping state with the crystal structure conformation of free Scc3^SA2^ (RMSD = 2.4 Å, ***Figure 1 – figure supplement 1C***) (*Hara et al., 2014; Shi et al., 2020*). In the gripping state, Scc3^Psc3^ interacts with the cohesin loader both along the N-terminal Scc2^Mis4^ handle, as well as the central Scc2^Mis4^ hook. Scc2^Mis4^ rearrangement into its extended form following ATP hydrolysis would disrupt at least some of these contacts, thereby terminating the close Scc3^Psc3^ - Scc2^Mis4^ juxtaposition (***Figure 1B***, right). A conformational change within the Scc2^Mis4^ handle is furthermore likely to weaken its interaction with the SMC hinge. We therefore hypothesize that, as a consequence of Scc2^Mis4^ structural rearrangements following ATP hydrolysis, the interaction between the Scc2-head module and the Scc3-hinge module resolves. While the Scc2-head module turns from the DNA gripping to its slipping state, the DNA binding characteristics of the Scc3-hinge module remain unaltered.

### Measured positional changes between the Scc3-hinge and Scc2-head modules

Above, we predicted positional changes of the Scc3-hinge module relative to the Scc2-head module, when comparing the gripping state with ATP post-hydrolysis states. To experimentally observe the relative positions of module components, we designed FRET reporters inserted at the hinge within Smc1^Psm1^, at the C-terminus of Scc3^Psc3^ and at the N-terminus of Scc2^Mis4(N191)^ (***Figure 2A***). Scc2^Mis4(N191)^ is an N-terminally truncated Scc2^Mis4^ variant missing the first 191 amino acids. The truncation abrogates the interaction with Scc4^Ssl3^, a factor important for *in vivo* cohesin loading onto chromatin. *In vitro*, using naked DNA as a substrate, Scc2^Mis4(N191)^ retains full biochemical capacity to promote gripping state formation, topological cohesin loading, as well as loop extrusion (*Chao et al., 2015; Higashi et al., 2020; Shi et al., 2020*). Based on our structural model, these reporter locations are within distances that should allow robust FRET signal detection in the gripping state. CLIP or SNAP tags, inserted at these positions, served as fluorophore receptors. We labeled these tags during protein purification with Dy547 and Alexa 647 dyes as donor and acceptor fluorophores, respectively (***Figure 2B*** and ***figure supplement 1A***). We then mixed labeled cohesin, either the Scc2^Mis4^-Scc4^Ssl3^ cohesin loader complex or labeled Scc2^Mis4(N191)^, a 3 kb circular double stranded plasmid DNA and ATP in the indicated combinations. To create the gripping state, we included all components but substituted ATP for the non-hydrolyzable nucleotide ground state mimetic ADP · BeF3-. Following Dy547 excitation, we measured the relative FRET efficiency, defined as the Alexa 647 intensity at its emission peak divided by the sum of Dy547 and Alexa 647 emission intensities.

**Figure 2.**
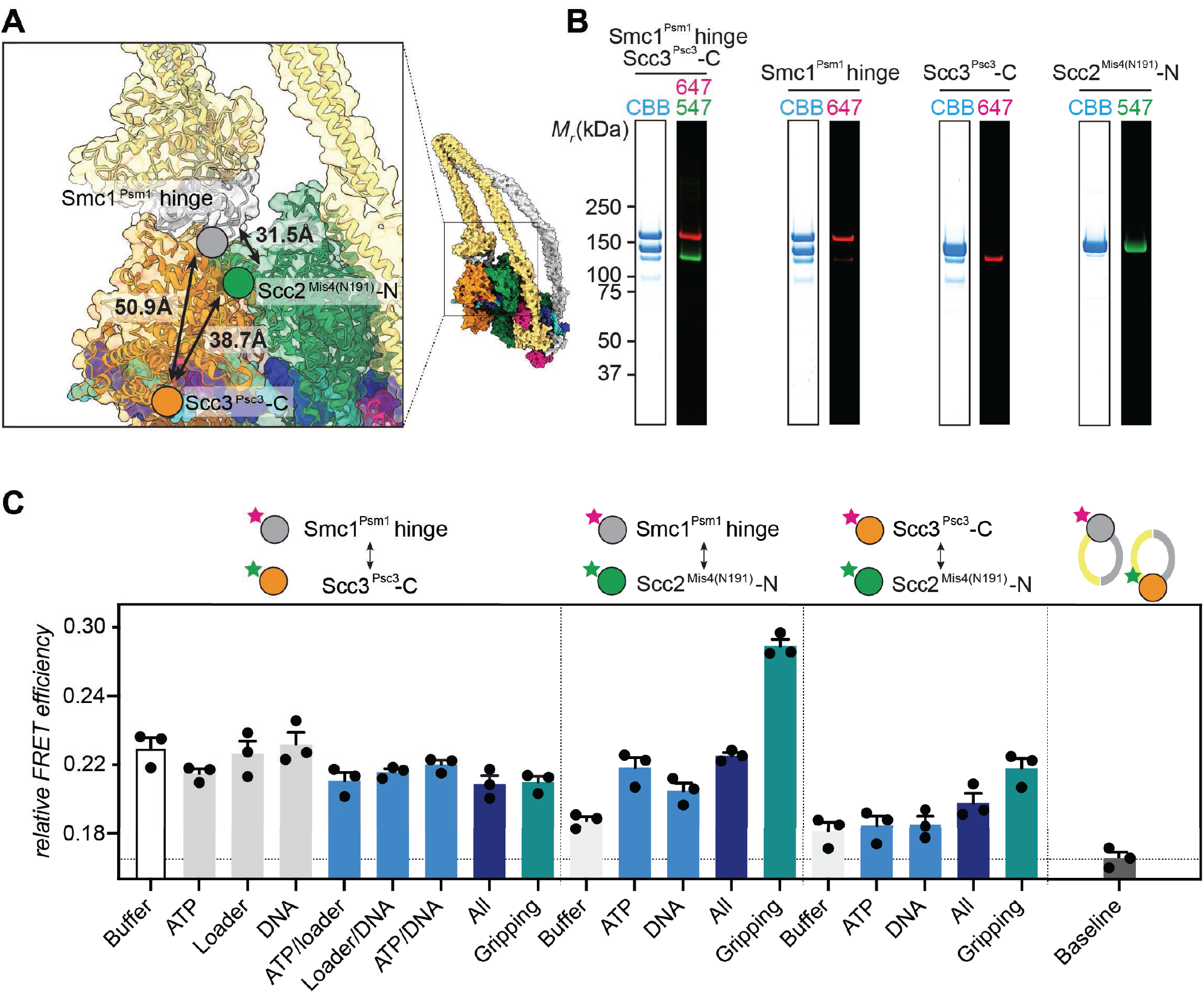
FRET-based conformational analysis of the Scc3-hinge and Scc2-head modules. **(A)** FRET reporter positions on the fission yeast cohesin structure in the DNA gripping state. The silver, orange and green circles mark the positions of the Smc1^Psm1^ hinge residue R593, the Scc3^Psc3^ C-terminal residue E959 and Scc2^Mis4^ residue P209, respectively. The Euclidean distances between the C_α_ atoms of these residues are indicated. **(B)** The purified and labeled cohesin complexes and Scc2^Mis4(N191)^ cohesin loader were analyzed by SDS-PAGE followed by Coomassie blue staining (CBB) or in gel fluorescence detection of the Cy547 (547) and Alexa 647 (647) dyes. **(C)** FRET efficiencies between the indicated elements were recorded under the conditions shown. The relative FRET efficiency is calculated as I_A_/(I_D_ + I_A_), where I_D_ is the donor emission intensity at 565 nm and I_A_ is the acceptor emission intensity at 665 nm resulting from donor excitation at 525 nm. The apparent FRET value observed using a mixture of single-labeled cohesins is indicated as a baseline. Results from three independent repeats of the experiments, their means and standard deviations are shown.

We first recorded FRET between the fluorophore pair at the Smc1^Psm1^ hinge and the Scc3^Psc3^ C-terminus. The FRET efficiency measured with the cohesin complex alone was 0.22 and displayed only negligible changes following the addition of one or more of the different cofactors. Even under conditions of gripping state formation, the FRET efficiency remained largely unchanged (***Figure 2C***). As a control, we prepared a mixture of singly Smc1^Psm1^ hinge and singly Scc3^Psc3^ C-terminus labeled cohesin complexes with the same concentrations of donor and acceptor dyes. This mixture provides a baseline for the apparent background FRET value due to spectral overlap. At 0.17 the measurement remained substantially below the FRET values observed when both fluorophores were incorporated within the same cohesin complex. This observation supports the idea that the SMC hinge and Scc3^Psc3^ lie in proximity of each other to form an Scc3-hinge module, consistent also with a biochemically observed Scc3^Psc3^-hinge interaction (*Murayama and Uhlmann, 2015*). Module formation was observed under all tested conditions, irrespective of the stage during cohesin’s ATP binding and hydrolysis cycle.

Next, we investigated the positioning of the Scc3-hinge module relative to the Scc2-head module. We measured FRET between a donor fluorophore at the Scc2^Mis4(N191)^ N-terminus and acceptor fluorophores at either the Smc1^Psm1^ hinge or at the Scc3^Psc3^ C-terminus. In both cases, we observed FRET at relatively low values under most conditions. Strikingly, the FRET efficiency of both fluorophore pairs markedly increased under conditions of gripping state formation (***Figure 2C***). The greatest FRET value of over 0.27 was recorded in the case of the Scc2^Mis4(N191)^ and Smc1^Psm1^ hinge pair in the gripping state. This is consistent with the shortest expected Euclidean distance of this fluorophore pair, amongst those tested. These observations confirm that the Scc3-hinge and Scc2-head modules come into proximity in the ATP-bound gripping state, as seen in the cryo-EM structures. The observations further suggest that this proximity is unique to the gripping state and that the two modules separate from each other at other times.

While the FRET efficiency between the Scc3-hinge and Scc2-head modules was low in conditions other than the gripping state, the measured values remained distinctly above background levels. Residual FRET could stem from a conformation in which the two modules remain within the FRET range of up to 100 Å. Alternatively, residual FRET could originate from transient encounters of the two modules that are free to dynamically move relative to each other, when not in the gripping state.

When using Scc2^Mis4(N191)^ as the FRET donor, its transitory interaction with cohesin becomes a confounding factor. Higher FRET efficiency in the gripping state could have been due to increased cohesin – Scc2^Mis4(N191)^ complex formation, rather than a conformational change. To exclude this possibility, we monitored the cohesin – Scc2^Mis4(N191)^ interaction by co-immunoprecipitation. This revealed equal interaction efficiencies under all of our incubation conditions (***Figure 2 – figure supplement 1B***). Therefore, the observed FRET differences cannot be explained by different cohesin – Scc2^Mis4(N191)^ complex stabilities. Rather, the FRET changes point to conformational transitions within the cohesin complex.

### The role of the Scc3-hinge module during topological DNA entry

What are the consequences of the Scc3-hinge module, and its movement relative to the Scc2-head module, for the DNA path during topological entry? Our earlier results suggested that DNA arrives from the top of the ATPase heads and usually passes the kleisin N-gate before reaching the gripping state (***Figure 3A***, panel ***a***) (*Higashi et al., 2020*). The kleisin N-gate initially opens as the consequence of ATP-dependent SMC head engagement (*Muir et al., 2020*). A positively charged kleisin N-tail then guides DNA through this gate *en route* to the gripping state. In this state, electrostatic contacts of the DNA, together with the Scc2^Mis4^ cohesin loader, shut the gate. In this configuration, the Scc3-hinge module docks onto the Scc2-head module. The straight DNA path through these two DNA binding elements in turn requires that a bend forms where the DNA arrives between the Smc1^Psm1^ and Smc3 ^Psm3^ coiled coils. The DNA path shown in ***Figure 3A***, panel ***a***, highlights the position of such a bend, based on our DNA-protein crosslink mass spectrometry data (*Higashi et al., 2020*). The notion of DNA bending in the gripping state finds further support from experiments where, using a magnetic tweezer setup, condensin binding to DNA under gripping state conditions has been observed to introduce a discrete DNA shortening step (*Ryu et al., 2020c*).

**Figure 3.**
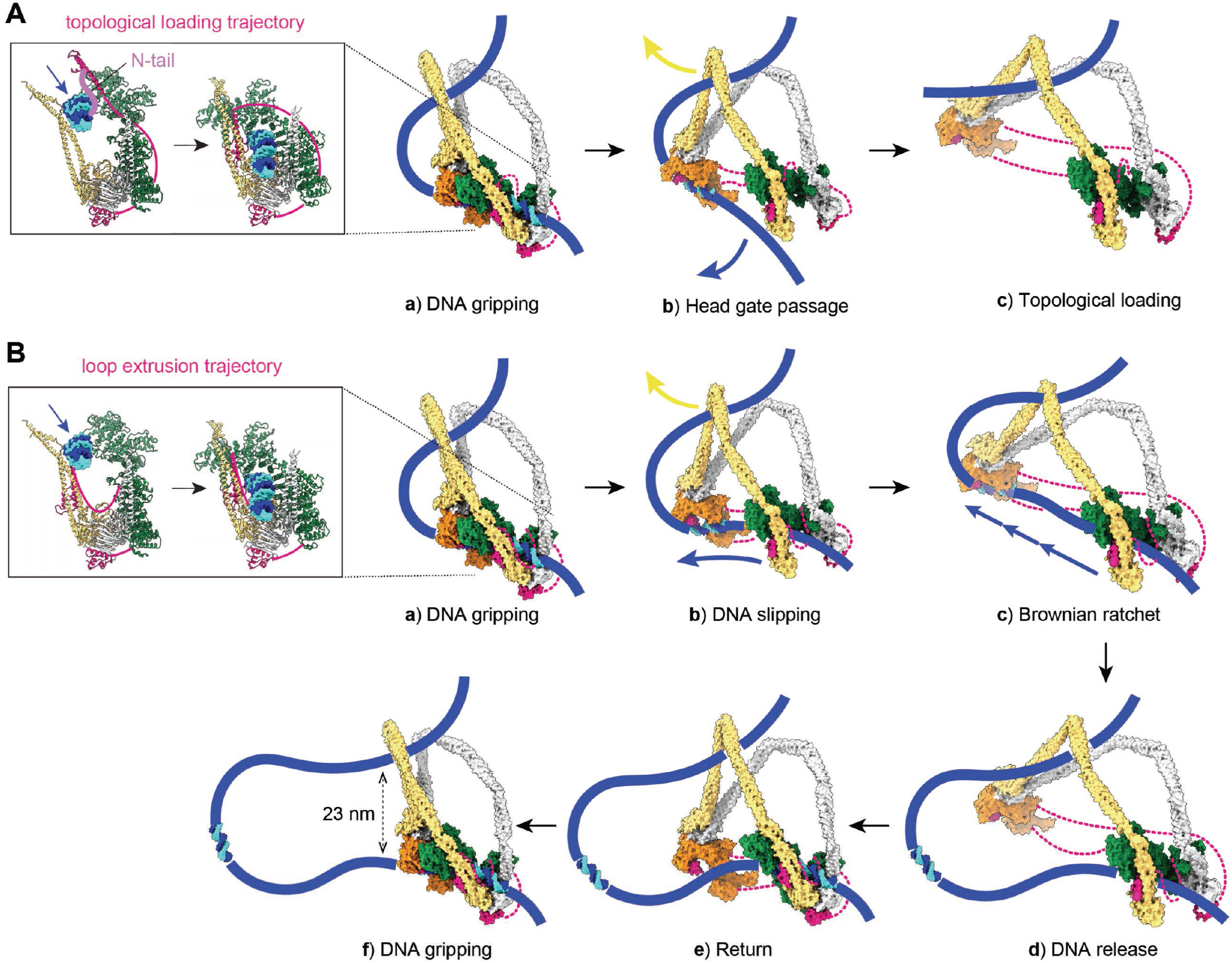
Molecular models for topological DNA entry into the cohesin ring and for loop extrusion. **(A)** Molecular model for topological DNA entry into the cohesin ring. ***a***, DNA enters through the open kleisin-N gate. DNA binding by both the Scc2-head and Scc3-hinge modules introduces a DNA bend. ***b***, ATP hydrolysis opens the SMC head gate. The swinging motion of the released Scc3-hinge module steers the DNA through the head gate to complete topological entry into the cohesin ring. ***c***, DNA is topologically entrapped in the Smc3-Scc3-kleisin-N chamber of the cohesin ring. **(B)** Molecular model for loop extrusion. ***a***, DNA arrives in the gripping state without passing the kleisin N-gate. ***b***, ATP hydrolysis leads to SMC head gate opening, but the kleisin path prevents DNA passage. The swinging motion of the Scc3-hinge module instead turns the bent DNA into a loop, while DNA slips along the Scc2-head module. ***c***, The amount of loop growth depends on the stochastic Brownian motion of the Scc3-hinge module. ***d***, The low DNA affinity of the Scc3-hinge module results in DNA release. ***e***, As long as the cohesin loader remains present, the Scc3-hinge module returns to form a new DNA gripping state upon nucleotide binding. ***f***, The next loop extrusion cycle begins. The in- and outbound DNAs are constrained by cohesin in our model at a distance of ∼ 23 nm, distinctly below the diameter of an unfolded cohesin ring and in line with recent measurements of the neck size of condensin when engaged in loop extrusion (*Ryu et al., 2020b*).

A stable DNA gripping state forms only if ATP hydrolysis is prevented by ATPase mutations or in the presence of non-hydrolyzable ATP. Usually, gripping state formation triggers ATP hydrolysis, resulting in ATPase head gate opening and Scc3-hinge and Scc2-head module uncoupling. This will initiate a swinging motion of the Scc3-hinge module and proximal coiled coil, with a pivot point at the elbow (***Figure 3A***, panel ***b***). No force is required to be transmitted along the SMC coiled coil. Rather, Brownian motion can take the Scc3-hinge module only in one direction, away from the Scc2-head module from which it was released. When we consider the consequence of the Scc3-hinge swinging motion on the DNA path, we make two observations. Firstly, the bent DNA straightens, an effect that in fact will energize the swinging motion. Secondly, the movement effectively steers the DNA through the ATPase head gate to complete topological entry into the cohesin ring.

Following ATPase head gate passage, DNA retains association with the Scc3-hinge module only for a limited time. DNA affinity to Scc3^Psc3^ and the kleisin middle region in isolation has been measured at around 2 *μ*M (*Li et al., 2018*), a relatively low affinity that implies a fast DNA off-rate once Scc3^Psc3^ has left the gripping state. DNA consequently finds itself in a cohesin chamber delineated by the Smc3^Psm3^ coiled coil, Scc3^Psc3^, as well as the unstructured part of the kleisin between the kleisin middle region that binds Scc3^Psc3^ and the kleisin N-gate (***Figure 3A***, panel ***c***). We refer to this space as cohesin’s Smc3-Scc3-kleisin-N chamber. Two separase recognition sites in Scc1^Rad21^, whose cleavage liberates DNA from the cohesin ring to trigger anaphase (*Tomonaga et al., 2000; Uhlmann et al., 2000*), are situated within this part of the kleisin unstructured region (***Figure 3 – figure supplement 1A***).

Single molecule imaging of topologically loaded cohesin on DNA showed that its diffusion is blocked by obstacles smaller than those expected to be accommodated by cohesin’s SMC compartment (*Stigler et al., 2016*). This observation is consistent with the possibility that DNA resides in a sub-chamber of the cohesin ring following topological loading. How durable the Scc3^Psc3^-hinge association is, whether DNA permanently resides inside the Smc3-Scc3-kleisin-N chamber, or whether subunit rearrangements take place following successful topological loading, *e*.*g*. when the cohesin loader is replaced by Pds5, remains to be further ascertained.

### An alternative gripping state that initiates and drives loop extrusion

The structured components of the gripping state do not by themselves contain information about whether DNA has in fact passed the kleisin N-gate. While mechanisms are in place to ensure kleisin N-gate passage, *e*.*g*. the kleisin N-tail, DNA might under certain conditions reach the gripping state without having passed this gate (***Figure 3B***, panel ***a***). What will be the consequence of ATP hydrolysis in such an alternative gripping state? ATPase head dissociation turns the Scc2-head module into its DNA slipping state, but the kleisin path prevents DNA from passing between the ATPase heads. The Scc3-hinge module again uncouples from the Scc2-head module, but now its diffusion-driven swinging motion cannot steer DNA through the head gate. The only way for the Scc3-hinge module to launch its swinging motion is to further bend the DNA, turning it into a loop, while DNA slips along the Scc2-head module (***Figure 3B***, panel ***b***). Maturation of the DNA bend in the gripping state into a DNA loop is likely to be energetically costly. Brownian motion will have to surpass the required activation energy, possibly after a certain delay (***Figure 3B***, panel ***c***).

Once a DNA loop is initiated, the extent of loop growth is limited by how far the Scc3-hinge and Scc2-head modules can separate from each other. The maximum separation is likely dictated by the kleisin unstructured regions that link Scc3^Psc3^ to the Scc2-head module. Their lengths of 135 amino acids (between the Scc2^Mis4^ and Scc3^Psc3^ binding sites) and 109 amino acids (between the Scc3^Psc3^ binding site and the kleisin C-terminal domain) gives a conservative estimate of ∼ 40 nm (*Ainavarapu et al., 2007*) (***Figure 3 – figure supplement 1B***). This distance allows considerable, but perhaps not complete, opening of the ∼ 47 nm long SMC proteins. As we will show below, the actual amount of loop growth is likely less and depends on the energy balance between DNA and cohesin that is reached by stochastic diffusive motion at the time when DNA dissociates (***Figure 3B***, panel ***d***).

After DNA dissociation from the Scc3-hinge module, there is a time without tight contact between the cohesin ring and the DNA loop. Thermal fluctuations lead to random loop size changes, depending on the probability of diffusion and on external forces that might apply to the DNA. As long as the Scc2^Mis4^ cohesin loader remains part of the Scc2-head module, the local proximity of all components means that a return of the Scc3-hinge module (***Figure 3B***, panel ***e***) and the establishment of a new DNA gripping state following nucleotide binding (***Figure 3B***, panel ***f***) are very likely events. The next loop extrusion cycle begins.

### A computational model for Brownian ratchet-driven DNA loop extrusion

In the above molecular model, two DNA binding modules within the cohesin complex generate a Brownian ratchet. The ratchet is operated by repeated cycles of ATP dependent DNA gripping state formation and its unidirectional dissolution following ATP hydrolysis. To evaluate whether such a mechanism is physically plausible and can indeed drive DNA loop extrusion, we constructed a structure-based molecular-mechanical model of the cohesin-DNA interaction and carried out computational simulations to explore its behavior. We modeled DNA as a discrete stretchable, shearable wormlike chain (dssWLC), which describes DNA at any arbitrary level of discretization with persistence length as the only parameter (***Figure 4A***) (*Koslover and Spakowitz, 2014*). We assumed the persistence length to be *Lp* = 50 nm (*Wang et al., 1997; Bustamante et al., 2000*). The cohesin coiled coil segments as well as the linkage between the two SMC heads were modeled using the same approach. Each coiled coil was represented as three beads that interact via a dssWLC (***Figure 4B***). This again leaves persistence length as the sole parameter that we chose such that it leads to a head-to-hinge distance distribution that matches experimentally measured head-to-hinge distances in a freely diffusing eukaryotic SMC complex (*Ryu et al., 2020b*).

**Figure 4.**
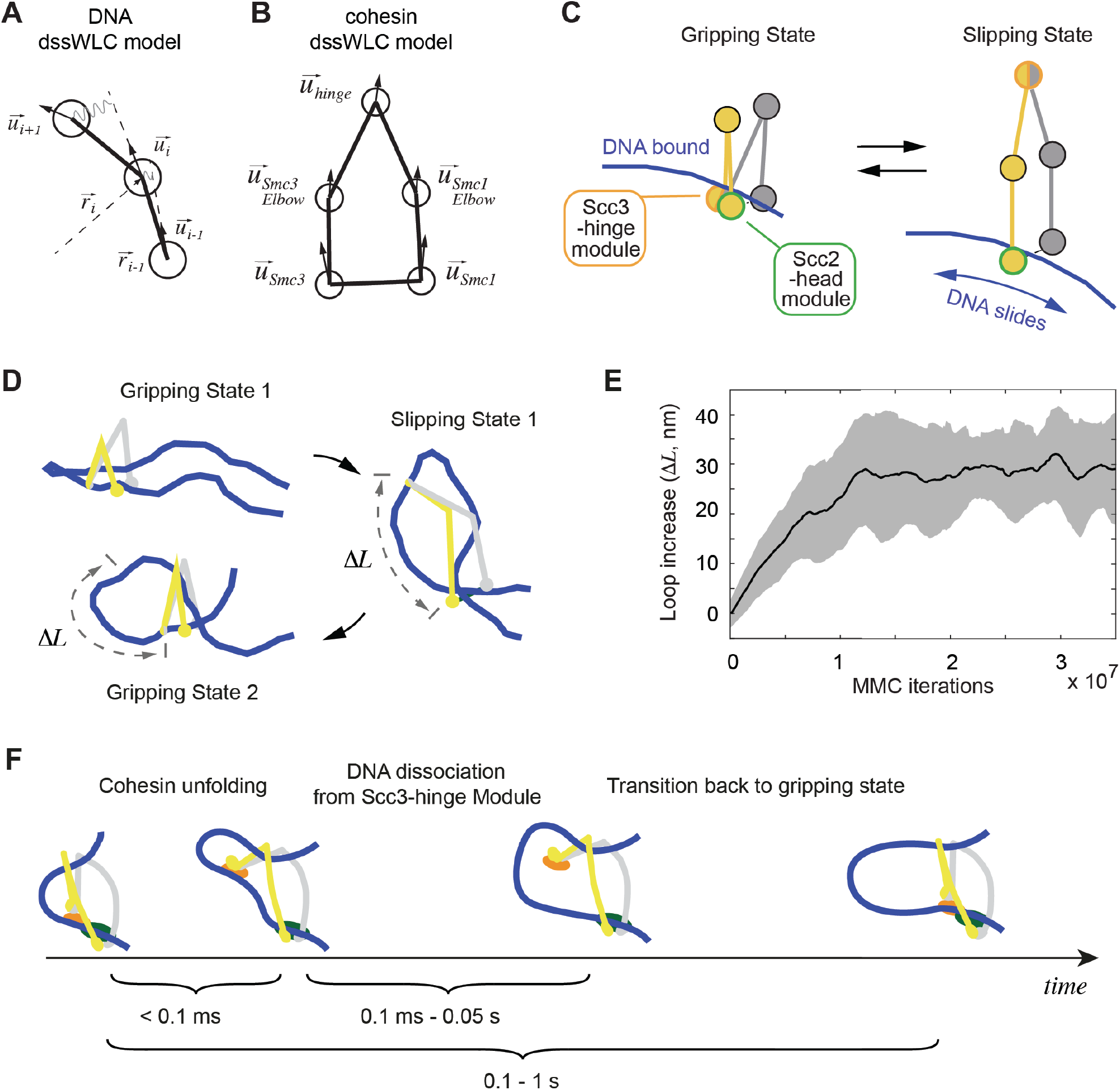
3D Metropolis Monte-Carlo simulation of Cohesin-DNA interactions. **(A)** Representation of DNA as a dssWLC model. DNA is split into 5 nm long segments. Each segment is described by its radius vector 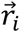 and a unit vector 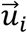 that defines segment orientation. **(B)** Cohesin is described by five beads corresponding to Smc1 and Smc3 heads and elbows as well as the hinge. The bead positions are defined by the corresponding vector radii (not shown) and their orientation by unit vectors. **(C)** The two equilibrium conformations of the model, corresponding to the gripping and slipping states. **(D)** Snapshots from a simulation started in a gripping state. The system was sampled 2.5·10^6^ times and then the equilibrium state was changed to the slipping state. It was sampled for another 8·10^6^ iterations, which led to cohesin unfolding and loop extension. At iteration 1 ·10^7^ the equilibrium conformation was returned to gripping state. **(E)** Loop length increase shown as function of Metropolis Monte-Carlo (MMC) iterations following the gripping to the slipping state transition. Before iteration zero, the system was equilibrated for 2·10^6^ rounds in the gripping state. The black line represents the average and the grey area the standard deviation across ten independent simulation replicates. **(F)** Schematic to show time progression of the DNA interaction with the Scc3-hinge module. Time intervals show indicative ranges of model parameters.

Based on our cryo-EM structure and biochemical observations, we define two states of the cohesin complex. In the gripping state, the Scc3-hinge and Scc2-head modules are engaged and the coiled coil elbows are folded. Both modules make stable contact with DNA (***Figure 4C***, Gripping State). In the second state, the slipping state, the Scc3-hinge and Scc2-head modules do not interact, allowing an unfolded cohesin conformation. In this state, the Scc2-head module permits free transverse DNA motion. DNA association with the Scc3-hinge module is defined by its equilibrium dissociation constant, as discussed below (***Figure 4C***, Slipping State).

First, we explored the dynamics of the transition between cohesin’s gripping and slipping states. As a starting point we assume that bent DNA is inserted into the cohesin ring in the gripping state. The first panel in ***Figure 4D*** shows a snapshot of this state after equilibration of the Metropolis Monte-Carlo algorithm (see Material and methods). We then simulated a transition to the slipping state. To this end, we disconnect the Scc3-hinge from the Scc2-head module and switch the Scc2-head module to its slipping state, while the Scc3-hinge module remains bound to DNA. We then sampled conformations until a new equilibrium was reached. As cohesin unfolds, DNA binding to the Scc3-hinge module limits DNA movement at the Scc2-head module to only one direction, towards an increased loop size (***Figure 4D***). The average increase in loop size during multiple repeats of this transition is ∼ 30 nm (***Figure 4E***). When we then allow DNA to detach from the Scc3-hinge module and switch cohesin back to the gripping state, the system readily resets and primes itself for the next cycle (***Figure 4D***). Our simulations reveal that repeated rounds of the states: “gripping -> slipping -> DNA detachment from the Scc3-hinge module -> gripping” results in continuous extrusion of DNA with an average loop size increase of ∼ 30 nm per cycle (***Supplementary Video 1***).

### DNA affinity of the Scc3-hinge module controls loop extrusion

For the above mechanism to achieve processive cycles of loop extrusion, we have to make two assumptions about the DNA contact at the Scc3-hinge module. First, DNA binding must persist for long enough, following the gripping to slipping state transition, to ensure biased DNA diffusion towards loop growth while cohesin unfolds (***Figure 4F***). An upper limit for the time it takes cohesin to unfold is given by the time of a diffusive process that separates the Scc3-hinge and Scc2-head modules. Assuming molecular masses of both modules in the 200 kDa range, it takes ∼ 0.1 ms for them to diffuse ∼ 50 nm apart (see Materials and methods). This time is an upper estimate. If cohesin opening was driven not merely by diffusion, but assisted by internal stiffnesses of the coiled-coils, this could speed up opening. Based on this estimate, our model predicts that the off-rate of DNA at the Scc3-hinge module should be slower than 1/0.1 ms = 10,000 s^-1^ in order for DNA to maintain Scc3-hinge module association until cohesin fully unfolds.

After cohesin has opened, two further scenarios are possible. Firstly, DNA could dissociate from the Scc3-hinge module before cohesin transitions back into the next gripping state. In this case a net loop length gain is made and the ensuing gripping state primes cohesin for the next round of loop extrusion (***Figure 4F***). Alternatively, cohesin could switch back to the gripping state before DNA is released from the Scc3-hinge module. In this situation, DNA ends up in the same position as the previous gripping state and there is no net loop size gain, resulting in an unproductive cycle. Based on these considerations, loop extrusion in our model is most effective when the lifetime of the DNA to Scc3-hinge module contact is shorter than the time it takes cohesin to transition back into the next gripping state. Such a lifetime would ensure that most reaction cycles result in net loop growth.

The ATP-bound DNA gripping state is an unstable, transient state, so we can expect cohesin to spend the majority of its time in the post-hydrolysis slipping state. We can then approximate the lifetime of this state based on cohesin’s ATP hydrolysis rate. This rate has been measured with a lower limit of ∼ 2 s^-1^ (*Murayama and Uhlmann, 2014; Ganji et al., 2018; Davidson et al., 2019*). As two ATPs are coordinately hydrolyzed by cohesin, this equates to a cycle rate of ∼ 1 s^-1^. Thus, our model predicts that efficient loop extrusion is achieved when the off-rate of DNA from the Scc3-hinge module is in the range of 1 – 10,000 s^-1^. While we do not know the actual off-rate, the equilibrium dissociation constant *K*_*d*_ between DNA and Scc3 has been measured at ∼ 2 *μ*M (*Li et al., 2018*). Assuming an association rate *K*_*on*_ typical for biomolecular interactions of ∼ 10^7^ M^-1^ s^-1^ (*Howard, 2001*), we arrive at a corresponding off-rate *K*_*off*_ = *K*_*d*_ × *K*_*on*_ of ∼ 20 s^-1^. This value sits well within the range predicted to support processive loop extrusion. The number ensures processivity even if cohesins that are actively engaged in loop extrusion undergo conformational cycles and hydrolyses ATP at an up to twenty times faster rate compared to that measured in bulk solution experiments.

### Loop extrusion driven by biased DNA diffusion

Having established that transitions between cohesin’s gripping and slipping states can drive directional DNA movements, we explored how this mechanism compares to available experimental observations of loop extrusion at realistic time scales. To do this, we constructed a simplified model of loop development. We assume that both DNAs that enter and exit cohesin can randomly diffuse in and out of the ring with rates depending on the DNA diffusion coefficient *D*. We then use a Monte-Carlo algorithm to simulate DNA loop dynamics as a function of time.

If we adopt a diffusion coefficient of ∼ 1 *μ*m^2^/s, as measured for cohesin movements on DNA following topological loading (*Davidson et al., 2016; Stigler et al., 2016*), we see that both strands randomly diffuse back and forth, leading to stochastic loop size changes (***Figure 5A***). Within a few minutes, a typical time frame used to microscopically observe DNA loop extrusion, the DNA loop size changes over a range of several kb. However, these random movements do not show a preferred direction and cannot drive loop extrusion.

**Figure 5.**
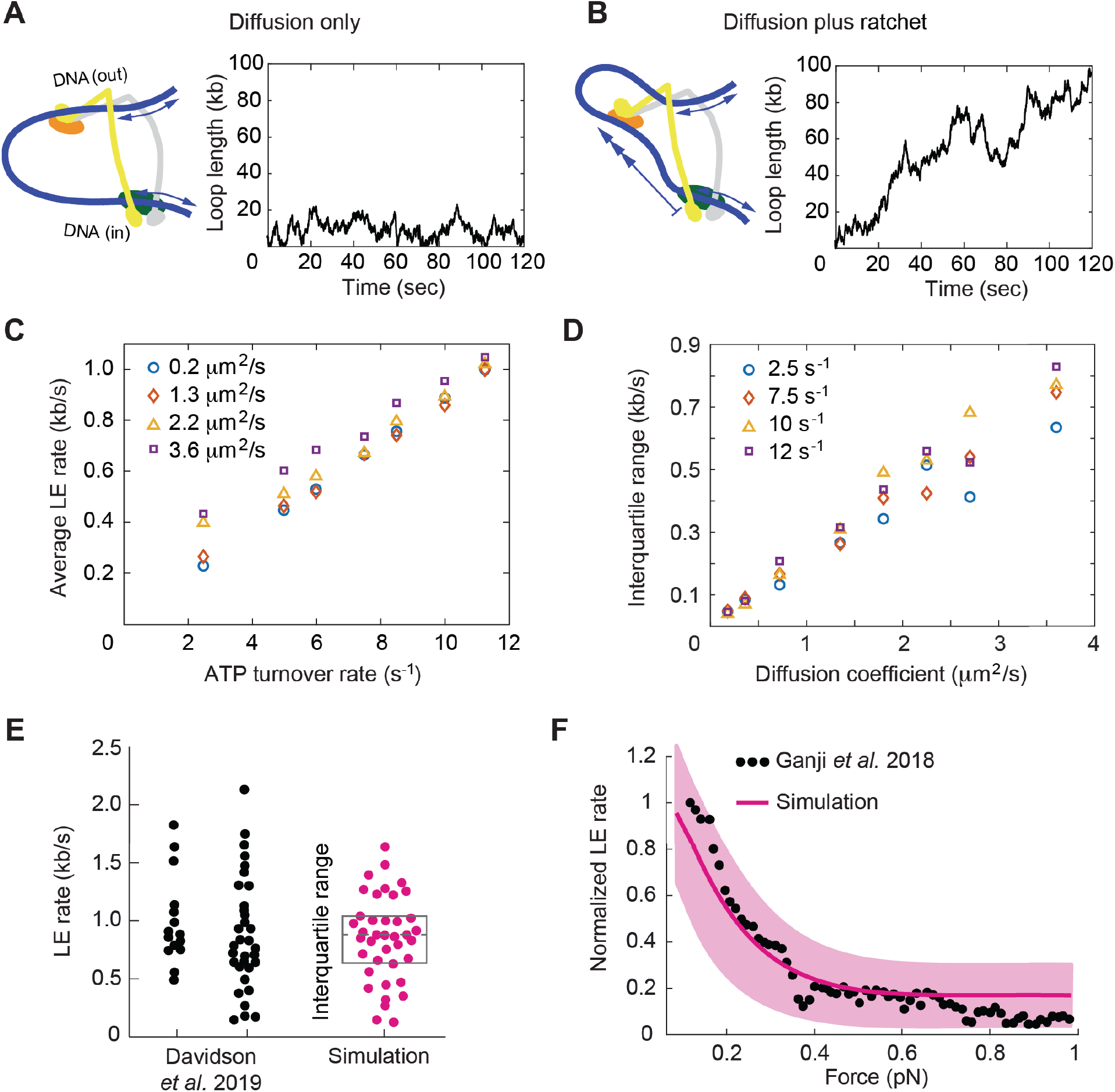
Loop extrusion by cohesin as a Brownian ratchet. **(A)** DNA loop size changes due to diffusion only of both DNAs. **(B)** DNA loop size changes due to random diffusion plus directed diffusion generated by cohesin’s gripping to slipping state transition at a rate of 9 s^-1^. **(C)** Simulated average loop extrusion (LE) rates as a function of the cohesin cycle (ATP turnover) rate. Symbol colors represent simulations with the indicated diffusion coefficients. **(D)** Scatter in the loop extrusion rates, generated during 3-minute simulations and quantified as the interquartile range, shown as a function of the outbound DNA diffusion coefficient. The diffusion coefficient for the inbound strand was 0.05 μm^2^/s. Symbol colors indicate simulations with different cohesin cycle rates. **(E)** Comparison of experimentally observed loop extrusion (LE) rate distributions of HeLa cell cohesin (left) and recombinant human cohesin (right) (*Davidson et al., 2019*) and an example of simulated data over the same time interval. **(F)** Comparison of the experimentally observed force dependence of condensin loop extrusion (LE) rates (G*anji et al., 2018*) with the simulated outcome. The pink line shows the median across 35 simulations performed with discrete force values at 0.1 pN increments. The light pink area shows the corresponding interquartile range. Parameters for the simulations in (**E**) and (**F**) were: cohesin cycle rate = 9 S^-1^, DNA diffusion coefficient =1.5 μm^2^/s, simulation time = 3 minutes.

This situation changes when the Scc3-hinge module engages with DNA in the gripping state and disengages predominantly in the slipping state. The Scc3-hinge module restricts DNA diffusion at the Scc2-head module to only one direction – towards loop growth. This effect applies only to the DNA that enters the loop at the Scc2-head module, but not to the DNA that exits cohesin. We assume that the latter DNA continues to diffuse randomly in both directions irrespective of the cohesin state. If we simulate directed DNA motion at the Scc2-head module of 30 nm per cohesin turnover cycle, we see how, overlaid over stochastic diffusive loop size fluctuations, the loop length steadily increases over time (***Figure 5B***).

We next explored how the variables in this model affect the outcome of loop extrusion. There are three independent variables: the two diffusion coefficients that describe the in and outward movements of the two DNAs that enter and exit cohesin, as well as the ATPase turnover rate, *i*.*e*. the lifetime of each slipping state. We simulated multiple 10-minute intervals of cohesin-DNA dynamics and compared loop extrusion rates extracted from these simulations to those determined in experiments (*Ganji et al., 2018; Davidson et al., 2019*).

These simulations revealed that the average loop extrusion rate is unaffected by the DNA diffusion coefficient (***Figure 5C***). Indeed, thermal movement of DNA has no net direction and therefore should not contribute to directed loop growth. Instead, the average loop extrusion rate *v* is simply a product of the step size during cohesin’s state transitions *L* and the frequency *γ* of these events:

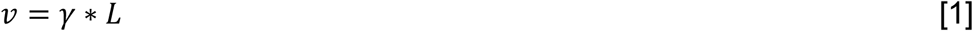

Using the value of *L* = 30 nm = 0.088 kb, we find that there must be at least 9 successful cohesin state transitions per second to reach experimentally observed average loop extrusion speeds of ∼ 0.8 kb s^-1^. The required rate of cohesin state transitions necessitates an equal rate of ATPase cycles. This means that a cohesin complex that is actively engaged in loop extrusion hydrolyzes ATP ∼ 9 times faster than average bulk solution ATP hydrolysis rates suggested.

A striking feature of experimentally observed loop extrusion is a high variation in loop growth rates (*Ganji et al., 2018; Davidson et al., 2019*). To obtain insight into the origin of these variations, we compared scatter in our modeled traces with experimental data. We quantified the scatter as the interquartile range, *i*.*e*. the range that contains 50% of datapoints around the median. This analysis revealed that extrusion rate variations strongly correlated with the DNA diffusion coefficient. The bigger the diffusion coefficient, the greater is the variation (***Figure 5D***).

A DNA loop consists of two DNAs that pass through cohesin and additional simulations showed that only the DNA with the higher diffusion coefficient determines the amount of scatter in extrusion speed (***Figure 5 – figure supplement 1***). A diffusion coefficient of ∼ 1.5 µm^2^/s resulted in a good match to the experimentally observed variation (***Figure 5E***). This diffusion coefficient matches the upper range of experimentally measured values for cohesin that is topologically bound to DNA (*Davidson et al., 2016; Stigler et al., 2016*). We imagine that the outward pointing DNA, which is not constrained by the Scc2-head module, shows the greater diffusion coefficient amongst the two DNAs. It might not interact strongly with cohesin and be largely free to diffuse. In addition to the high variations of loop extrusion rates, the low friction at the outward pointing DNA could also explain why DNA can be readily pulled from a condensin complex undergoing loop extrusion (*Kim et al., 2020*).

Finally, we explored how Brownian ratchet-driven loop extrusion in our model is affected by external force. Cohesin unfolding in the slipping state, which underlies biased DNA diffusion, is driven by thermal motion. Its rate in response to external force is given by:

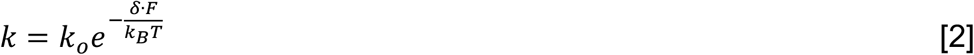

Where *F* is the external force and *δ* = 30 nm from our simulations (*Howard, 2001*). Introducing this dependency into our model, we find good agreement between simulations in the presence of a range of applied external forces and the experimentally observed decay of the force-velocity relationship (***Figure 5F***) (*Ganji et al., 2018*). The similarity between the simulated and experimentally observed responses to external force, as well as the high variation of experimentally observed loop extrusion rates, support the idea of a diffusion driven molecular mechanism of loop extrusion.

### Symmetric loop extrusion as a special case of asymmetric loop extrusion

In our molecular model of DNA loop extrusion, the Brownian ratchet acts only on the DNA that enters the cohesin ring through the Scc2-head module. No directional effect is exerted on the DNA that exits cohesin. This results in asymmetric loop extrusion (***Figure 6A***, Asymmetric loop extrusion), a scenario that is seen in the case of the condensin complex (*Ganji et al., 2018; Golfier et al., 2020*). In contrast, the experimental observations suggest that the cohesin complex symmetrically extrudes DNA loops (*Davidson et al., 2019; Kim et al., 2019*). How could this difference be explained?

**Figure 6.**
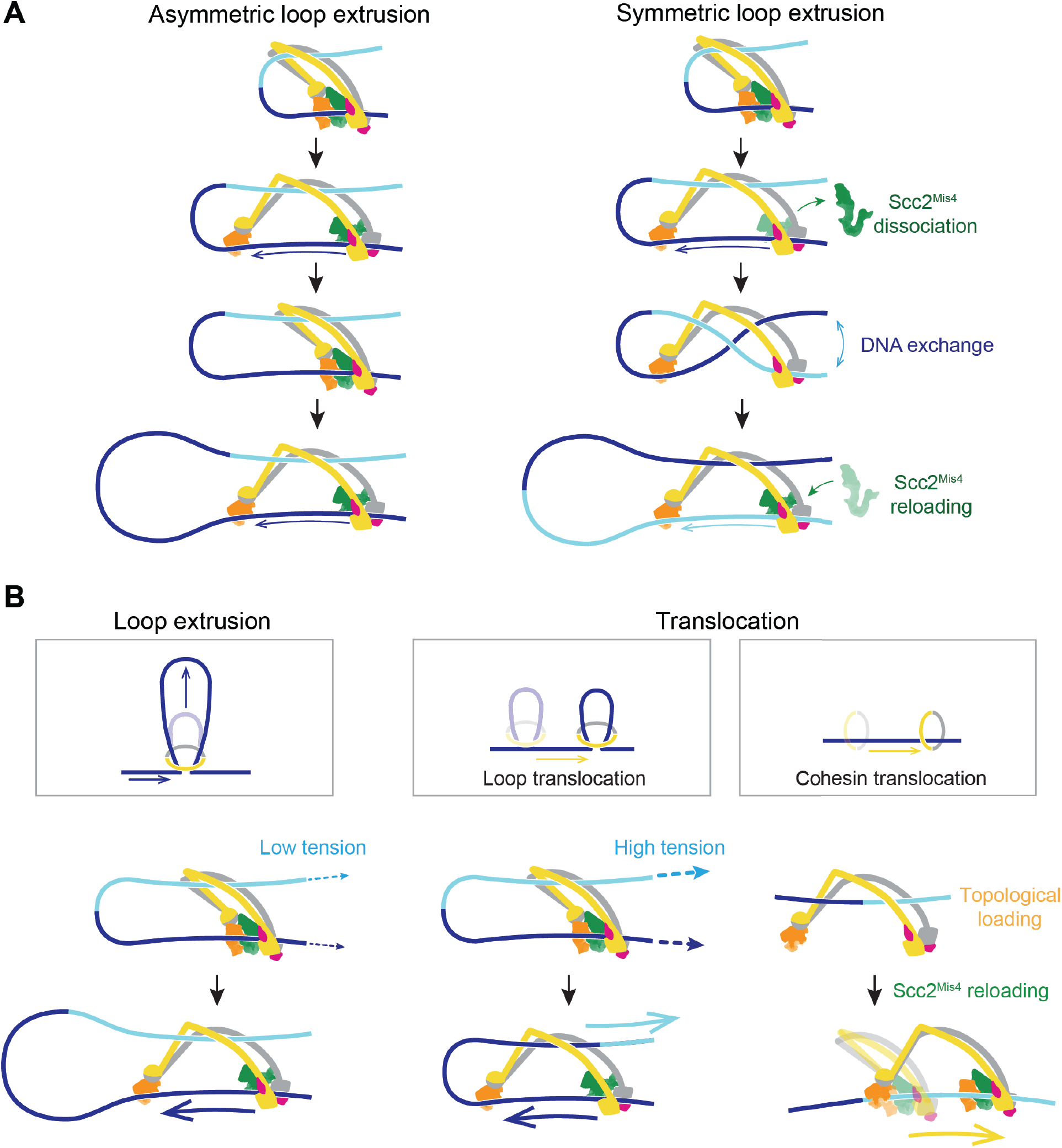
A model for asymmetric and symmetric loop extrusion, as well as possible mechanisms for cohesin translocation along stretched DNA. (**A**) A model for symmetric loop extrusion as a special case of asymmetric loop extrusion. Continued action of the Brownian ratchet on the DNA that enters the cohesin ring results in *Asymmetric loop extrusion*. If the cohesin loader dissociates, the Brownian ratchet disassembles. Reassembly of the Brownian ratchet following Scc2^Mis4^ reloading gives both DNAs an equal chance to become part of the ratchet. Over time, alternating asymmetric extrusion of both DNAs results in apparent *Symmetric loop extrusion*. (**B**) Two possible mechanisms for cohesin translocation along DNA under tension. *Loop extrusion* is possible only against very weak DNA counterforces. If loop growth stalls due to DNA tension, the continued operation of the Brownian ratchet leads to Loop *translocation*. Alternatively, cohesin might at first topologically load onto DNA. Recurring gripping state formation and resolution following topological loading results in Brownian ratchet driven *Cohesin translocation*.

In our model, the cohesin loader is a stable part of the cohesin complex, confining the DNA that enters the ring. However, Scc2^Mis4^ is not a permanent component of the cohesin complex and we can expect this subunit to turn over with a certain frequency. Suggestive of turnover, the continuous presence of cohesin loader in the incubation buffer is a requirement for processive loop extrusion by human cohesin (*Davidson et al., 2019*). If we picture a situation in which Scc2^Mis4^ dissociates from the cohesin complex, DNA will be released from the Scc2-head module (***Figure 6A***, Symmetric loop extrusion). The DNAs that enter and exit the cohesin ring are now indistinguishable and, once Scc2^Mis4^ rebinds, both DNAs have an equal chance to associate with the Scc2-head module during the new gripping state formation. Every round of cohesin loader dissociation and reloading thereby results in a one-in-two chance that the extruded DNA strand will switch. Averaged over time, this takes the appearance of symmetric loop extrusion.

### Loop extrusion versus directional cohesin translocation

In addition to loop extrusion, single molecule studies have reported ATP-dependent unidirectional translocation of cohesin and condensin along DNA (*Terakawa et al., 2017; Davidson et al., 2019*). The experimental conditions under which both complexes move along DNA, or extrude DNA loops, are largely similar. A difference lies in the DNA substrates on which translocation was observed. These substrates were stretched, either by liquid flow or by being doubly tethered to a flow cell surface. We have seen above that a Brownian ratchet is able to extrude a DNA loop only against very small externally applied forces (***Figure 6B***, Loop extrusion). If DNA is longitudinally stretched, loop extrusion will be limited to a small loop size. As the Brownian ratchet continues to deliver DNA to enlarge the loop, the stretching force begins to extract the DNA from the opposite side (***Figure 6B***, Loop translocation). Instead of promoting loop growth, the Brownian ratchet now fuels a motor that moves along the DNA. Consistent with this outlined scenario, a small DNA loop was observed to precede initiation of condensin translocation along stretched DNA (*Ganji et al., 2018*).

We can imagine an alternative scenario by which cohesin could turn into a Brownian ratchet-driven motor. Following successful topological loading onto DNA, cohesin might be able to return to a gripping state if the cohesin loader returns (***Figure 6B***, Cohesin translocation), *e*.*g*. if Scc2^Mis4^ is not replaced by Pds5. This possibility finds support from the observation that the cohesin loader retains the ability to stimulate cohesin’s ATPase following completion of topological loading (*Çamdere et al., 2015*). If the gripping state now reforms without DNA passing the kleisin N-gate, the following post-hydrolysis state in which the Scc2-head module turns to its slipping state, while the Scc3-hinge module initiates directed Brownian motion, leads to cohesin translocation along DNA. This second model for directed cohesin movement is not mutually exclusive with the earlier described ‘loop translocation’ model. Both models make the prediction that, similar to what is observed during loop extrusion, cohesin is a weak translocating motor that can be stalled by very small forces.

### Possible outcomes of cohesin collisions with DNA-bound obstacles

DNA loop extrusion by cohesin and condensin has so far only been observed *in vitro* and only using naked DNA substrates. *In vivo*, DNA is densely decorated by histones and other DNA binding proteins related to transcription, DNA replication and other forms of DNA metabolism. If we portray a loop extruding cohesin complex in its slipping state next to a nucleosome (***Figure 7A***, left), it becomes apparent that a nucleosome is too big to pass through the channel formed between Scc2^Mis4^ and the SMC ATPase heads. However, a possible path for nucleosome bypass opens up when the Scc2^Mis4^ cohesin loader transiently dissociates. DNA now passes in and out of cohesin through the Smc3-Scc3-kleisin-N chamber (***Figure 7A***, Nucleosome bypass, and ***figure supplement 1A***). In the case of topologically loaded cohesin, this channel seems in principle wide enough to allow nucleosome bypass, albeit denser nucleosome arrays block purely diffusive cohesin movement (*Stigler et al., 2016*). During loop extrusion, on one hand, Brownian ratchet-driven directional cohesin movement will facilitate nucleosome bypass. On the other hand, only one DNA lies in the Smc3-Scc3-kleisin-N chamber following topological loading, but both in and outward pointing DNAs must be accommodated during loop extrusion. If both DNAs are histone-bound, considerable steric constraints will substantially slow down loop extrusion. Compared to cohesin, loop extrusion by the condensin complex is likely to be equally, if not more, impeded by DNA-bound obstacles as the Scc2^Mis4^ analogous subunit, Ycs4^Cnd1^, is a stable condensin component (*Hirano et al., 1997; Lee et al., 2020*). While Ycs4^Cnd1^ might sporadically dissociate from the SMC heads to open up a bypass chamber, its persisting kleisin association will form an additional hindrance during obstacle bypass.

**Figure 7.**
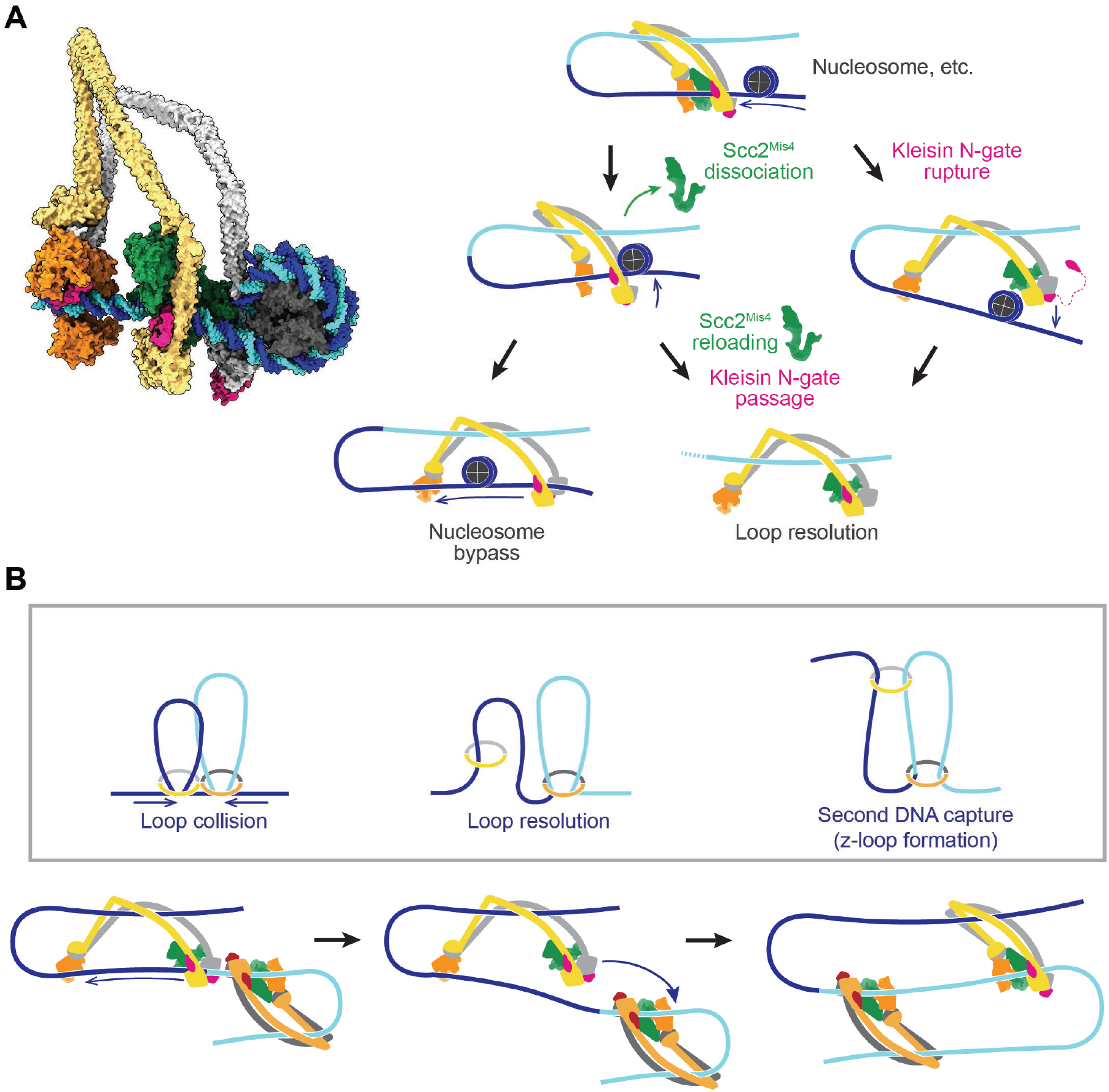
Possible outcomes of loop-extruding cohesin encounters with nucleosomes, or head-on encounters of two loop-extruding condensins. (**A**) Structural model of cohesin in the slipping state encountering a nucleosome (left). Three possible outcomes are shown on the right. If the cohesin loader dissociates, an Scc3-hinge module stroke can achieve *Nucleosome bypass* before the next gripping state forms. Alternatively, cohesin loader dissociation and reloading opens a renewed opportunity for *Kleisin N-gate passage* by DNA, resulting in *Loop resolution* and topological cohesin loading onto DNA. The impact of a nucleosome collision might also cause *Kleisin N-gate rupture*, which again results in *Loop resolution* and topological cohesin loading onto DNA. (**B**) A model for z-loop formation following head-on *Loop collision* of two loop extruding condensin complexes. One of the two condensins undergoes *Loop resolution*, by either kleisin N-gate passage or N-gate rupture. The now topologically bound condensin complex diffuses in search for a substrate for Second DNA capture, resulting in *z-loop formation*. During two-sided z-loop growth, depicted here, both in and outward pointing DNAs move through both condensin complexes. One is directed by each condensin’s Brownian ratchet, the other by the motion of the opposite condensin.

An alternative outcome of cohesin – nucleosome collisions is that loop extrusion at least temporarily stalls. Once the cohesin loader dissociates, DNA is released from the Scc2-head module and free to move within the Smc3-Scc3-kleisin-N chamber. As Scc2^Mis4^ rejoins the complex, there is a new chance for successful kleisin N-gate passage during gripping state formation, as originally foreseen during topological cohesin loading onto DNA (***Figure 7A***, Kleisin N-gate passage). If this occurs, the DNA loop resolves following ATP hydrolysis, resulting in cohesin topologically embracing one DNA.

There might be yet another possible outcome of cohesin-nucleosome encounters. Given the thrust of a diffusion-mediated collision between cohesin and an obstacle, the closed kleisin N-gate that prevented DNA passage through the SMC heads might rupture (***Figure 7A***, Kleisin N-gate rupture). If this happens, the extruded DNA loop will again resolve, resulting in cohesin topologically embracing one DNA. One can imagine that frequent stalling on a nucleosome-dense chromatin template prevents processive loop extrusion. The encountered obstacles could be seen as triggering a ‘proofreading’ mechanism, prompting recurring attempts at kleisin-N gate passage, eventually achieving topological loading onto DNA.

### Z-loop formation by SMC ring complexes

Chromosomal loading sites for cohesin complexes lie at considerable distances from each other. Nevertheless, we cannot outright dismiss the possibility that loop extruding cohesin complexes might collide in head-on encounters. *In vitro* observations of colliding, loop-extruding condensin complexes have revealed that the complexes are able to traverse each other to form distinctive z-loop structures (*Kim et al., 2020*). Can this behavior be explained by our molecular model of SMC complex function? Upon encounter, condensins have been observed to pause for a period of time. This is consistent with two Brownian ratchets that collide (***Figure 7B***, Loop collision). A way out of this conflict is provided by one of the two above mentioned loop resolution pathways, kleisin N-gate passage or N-gate rupture. Both allow one of the colliding condensins to resolve their loop and turn into a topologically loaded condensin (***Figure 7B***, Loop resolution). The newly gained freedom of movement of the singly tethered condensin allows it to diffuse. If, akin to how cohesin entraps a second single stranded DNA (*Murayama et al., 2018*), condensin now engages with a second double stranded DNA that lies beyond the colliding condensin complex, a z-loop is formed (***Figure 7B***, second DNA capture). Various outcomes have been observed during z-loop formation *in vitro*, including one- and two-sided z-loop growth. It is possible to explain both behaviors if we assume that second DNA capture during z-loop formation results in either topological capture of the second DNA or results in the second DNA entering loop extrusion mode.

## Discussion

Here we use structures of an ATP-bound DNA gripping state (*Higashi et al., 2020; Shi et al., 2020*) as the starting point to develop a molecular framework for DNA entry into the cohesin ring, as well as to show how a subtle but important difference in the starting topology turns this gripping state into a Brownian ratchet that can fuel DNA loop extrusion. The resulting model of cohesin function is able to explain numerous experimentally observed features of both topological loading and loop extrusion and makes a number of testable predictions.

### Use of the energy from ATP binding and hydrolysis during topological DNA entry

Any model of cohesin function must explain how the energy from ATP binding and hydrolysis is used to fuel or regulate its activities. ATP binding leads to SMC head dimerization and to a conformational change at the Smc3^Psm3^ neck that favors kleisin N-gate opening (*Higashi et al., 2020; Muir et al., 2020*). However, it appears that kleisin N-gate opening is not hard-wired and it is possible that head engagement occurs sometimes without kleisin N-gate opening (we will consider this possibility further, below). Dimerization importantly creates a composite DNA binding surface on top of the ATPase heads. The next steps in cohesin’s reaction cycle are now likely driven by the binding energy of DNA itself. First, DNA establishes contact with extensive positive surface charges on the ATPase heads. Next, the DNA engages Scc2^Mis4^, which turns into its gripping state conformation as its positively charged surface residues embrace the DNA. Together, the DNA and cohesin loader also establish contact with and thereby close the kleisin N-gate behind the DNA.

The DNA-induced Scc2^Mis4^ conformational change creates a docking interface for Scc3^Psc3^. The latter subunit joins the complex by simultaneously contacting the DNA, as well as by recruiting the SMC hinge. The binding energy released from these additional interactions compensates for the energetic cost of introducing a DNA bend, required to reach this configuration. Assembly of an energy-loaded gripping state is now complete and DNA has entered the cohesin ring through the kleisin N-gate. ATP hydrolysis now serves to dissolve the gripping state, however dissolution equates not merely to dispersal of the components. The geometric arrangement of the Scc3-hinge and Scc2-head modules in the gripping state, as well as their linkage by the folded coiled coils, mean that Brownian motion-driven dissolution generates a directed swinging movement of the Scc3-hinge module that promotes DNA head gate passage to complete topological DNA entry into the cohesin ring.

### Use of the energy from ATP binding and hydrolysis during loop extrusion

If DNA did not pass the kleisin N-gate, all other considerations for gripping state assembly remain unaltered. The binding energies of ATP and the DNA together again create an energy loaded state, the dissolution of which initiates a Brownian motion-driven swinging movement of the Scc3-hinge module. However, the kleisin prevents passage between the ATPase heads, instead resulting in DNA slippage along the Scc2-head module and the initiation of a DNA loop. The binding energy from ATP in this model operates a Brownian ratchet, while the movement of DNA is driven by thermal fluctuation. Processive loop extrusion by the Brownian ratchet depends on repeated gripping state formation and dissolution.

A key feature that determines the processivity of the Brownian ratchet is the half-life of the Scc3-hinge module interaction with DNA. This interaction should last long enough to guide directed diffusion but short enough so that DNA is released before the next gripping state forms. The corresponding component in the condensin complex is the putative Ycg1^Cnd3^-hinge module. Observations with condensin harboring a DNA binding site mutation in Ycg1^Cnd3^ revealed greatly compromised loop extrusion (*Ganji et al., 2018*), consistent with an important role of this element. To further evaluate our loop extrusion model, it will be important to modulate DNA affinity at both the Scc3-hinge and Scc2-head modules to explore their predicted effects on the efficiency of loop extrusion.

Biological motors typically couple ATP binding and hydrolysis-dependent conformational changes to their motion. This allows for robust movements in the presence of counteracting forces (*Howard, 2001*). We cannot exclude the possibility that cohesin also uses an ATP-dependent mechanism to control the relative positions of the Scc3-hinge and Scc2-head modules, *e*.*g*. SMC coiled coil stiffness could store torsional energy during gripping state formation. Release following ATP hydrolysis could then add a power stroke to loop extrusion by the Scc3-hinge module. However, SMC coiled coils appear to be very flexible (*Eeftens et al., 2016; Ryu et al., 2020b*) and as yet there is no evidence for energy coupling between the ATPase heads and hinge. Our computational simulations suggest that a purely diffusion driven Brownian ratchet recapitulates experimental observations of loop extrusion well.

### The importance of kleisin N-gate passage

A conserved kleisin N-tail interacts with DNA and ensures kleisin N-gate passage before the gate closes during gripping state formation (*Higashi et al., 2020*). Why then does kleisin N-gate passage sometimes fail, resulting in loop extrusion? A potentially relevant observation is that the biochemical reconstitution of topological cohesin loading onto DNA (*Murayama and Uhlmann, 2014*), as well as loop extrusion experiments (*Ganji et al., 2018*), are helped by unphysiologically low salt concentrations. Electrostatic interactions between DNA and cohesin, which characterize the gripping state, are stronger when less salt competes with them in the incubation buffer. While these enhanced DNA-protein contacts likely benefit the biochemically reconstituted reactions, the low ionic strength will also affect protein-protein interactions. For instance, electrostatic interactions contribute to keeping the kleisin N-gate closed and these are augmented in a low salt buffer. Therefore, the increased efficiency of gripping state formation at low salt concentrations might come at the cost of compromised kleisin N-gate operation. Whether the coupling of ATPase head engagement and kleisin N-gate opening is more stringent at physiological salt concentrations will be important to investigate, as will be ways that improve efficient cohesin function at physiological salt concentrations.

A hint for the importance of kleisin N-gate regulation comes from recent experiments performed to locate the DNA during its topological loading into the cohesin ring. Immediately upon cohesin addition to DNA, before topological loading becomes detectable, the DNA takes up a position in which chemical crosslinkers can trap it within what have been termed engaged-SMC and engaged-kleisin compartments (*Collier et al., 2020*). We imagine that this reflects the DNA approach between the SMC coiled coils, frequently passing through disengaged heads before the ATP-dependent series of topological loading events commences (***Figure 7 – figure supplement 1B***). Tight coupling of ATPase head engagement to kleisin N-gate opening will then be crucial to bring DNA onto the right path to topological entry into the cohesin ring.

### On the origin of loop extrusion

Could evolution have designed SMC complexes to be loop extruding Brownian ratchets? If the primordial function of SMC complexes was that of loop extruders, we should expect the loop extruding mechanism to be conserved in evolutionary ancient SMC complexes. Our model of loop extrusion suggests that cohesin’s DNA-interacting HEAT repeat subunits are key components of the Brownian ratchet that drives the process. These HEAT repeat subunits are relatively modern additions to SMC complexes. Evolutionarily older SMC complexes that are reflected in today’s prokaryotic SMC complexes, as well as in the Smc5-Smc6 complex, contain in place of HEAT subunits two smaller *k*leisin *i*nteracting *t*andem winged helix *e*lements (Kite) (*Palecek and Gruber, 2015*). While Kite subunits interact with DNA (*Zabrady et al., 2016*), they show important differences from how HEAT subunits are incorporated into SMC complexes. Kite subunits bind in close proximity to each other to a relatively short kleisin unstructured region (*Woo et al., 2009; Bürmann et al., 2013*). This observation makes it hard to imagine that a similar ratchet mechanism operates, which requires the DNA binding modules to separate from each other. The placement of Kite subunits within these SMC complexes might also change the way in which ATPase head engagement is coupled to kleisin N-gate opening. Further investigation of topological DNA entry into SMC-Kite complexes (*Kanno et al., 2015; Niki and Yano, 2016*), as well as their mechanism of gate operation, will provide important insight into the question whether these complexes can act as loop extruders.

### Implications for *in vivo* loop extrusion

Cohesin topologically entraps DNA to promote sister chromatid cohesion (*Haering et al., 2008*). It is possible that kleisin N-gate passage is an error-prone event and that accidental failure of kleisin N-gate passage during a topological DNA loading attempt initiates DNA looping. A close-by obstacle would soon stall loop extrusion and allow proofreading in the form of kleisin N-gate passage or N-gate rupture to reinstate topological loading. This scenario portrays loop extrusion as an unwanted, and possibly rare, side effect of topological cohesin loading. Alternatively, was it a cunning evolutionary twist that allowed cohesin to add loop extrusion to its repertoire of DNA acrobatics?

An obvious challenge to DNA loop extrusion *in vivo* is the presence of histones and other DNA binding proteins. Our model predicts that obstacle bypass is possible for loop extruding cohesins. At the same time, it suggests that obstacles reduce the speed and processivity of extrusion, especially when present at a high density on both the inward and outward directed DNA. Diffusion of topologically loaded cohesin along DNA is substantially slowed down by obstacles (*Davidson et al., 2016; Stigler et al., 2016*) and we expect that the same applies during loop extrusion. Recent studies reported DNA compaction of histone-bound DNA by cohesin and condensin, but whether these SMC complexes indeed bypassed histones in these experiments is not yet known (*Kim et al., 2019; Kong et al., 2020*). DNA loop extrusion was also observed in *Xenopus* egg extracts, both under interphase and mitotic conditions. However, loop extrusion was detectable only following histone depletion (*Golfier et al., 2020*). Obstacle encounters during *in vitro* loop extrusion are an obvious and important area for experimental investigation.

Many cellular DNA transactions make use of chromatin remodellers and histone chaperones to navigate the nucleosome landscape. Indeed, *in vitro* reconstituted chromosome assembly using purified condensin and histones depends on the histone chaperone FACT (*Shintomi et al., 2015*). While FACT facilitates DNA access by loosening the grip of histones on DNA, this histone chaperone does not possess catalytic activity or expend energy (*Zhou et al., 2020*). The energy for FACT-assisted nucleosome removal during transcription and DNA replication stems from the RNA or DNA polymerases that move along the DNA. How far FACT facilitates nucleosome eviction by a Brownian ratchet-fueled SMC complex will be interesting to investigate. In addition to FACT, one could imagine that ATP-dependent chromatin remodelers aid *in vivo* loop extrusion. When studying the contribution of histone chaperones and chromatin remodelers, we have to be aware that topological cohesin and condensin loading onto chromosomes also requires free DNA access that these enzymes provide (*Toselli-Mollereau et al., 2016; Garcia-Luis et al., 2019; Muñoz et al., 2019*).

Loop extrusion by SMC complexes is a captivating molecular event that provides an intuitive explanation for chromosome loop formation, explaining chromatin domains seen in chromosome conformation capture experiments (*Suhas et al., 2017*). However, DNA extrusion is not the only explanation for chromosome loop formation. Alternatively, cohesin and condensin could generate loops by sequentially topologically embracing two DNAs that come into proximity by Brownian motion, a mechanism that we refer to as diffusion capture (*Cheng et al., 2015; Gerguri et al., 2021*). Interactions between more than one cohesin binding site in the diffusion capture model could also arise from bridging-induced phase separation (*Ryu et al., 2020a*). When cohesin is depleted and re-supplied to human cells, small and large loops form with similar kinetics, a behavior that is more readily explained by a diffusion-mediated process than by gradual loop growth (*Suhas et al., 2017*).

In addition to cohesin and condensin, which constitute weak diffusion-driven motors, chromosomes harbor abundant, strong DNA translocases in the form of RNA polymerases. These translocases are known to push SMC complexes ahead as they move along chromosomes during gene transcription (*Lengronne et al., 2004; Davidson et al., 2016; Ocampo-Hafalla et al., 2016; Busslinger et al., 2017*). We have suggested that, following loop formation by diffusion capture, RNA polymerases provide extrinsic motor activity that promotes loop expansion (*Uhlmann, 2016*). Such transcription-dependent extrinsic loop expansion could explain chromatin domain features in an analogous fashion to cohesin moving on its own accord (*Bailey et al., 2020*). The role of RNA polymerase-dependent SMC complex movements in chromosome architecture deserves further attention.

### Outlook

Our molecular proposal for SMC complex function informs the evaluation how loop extrusion by SMC complexes might contribute to chromosome function. More importantly, while topological loading onto DNA and loop extrusion share many reaction steps, the two mechanisms also differ. The next challenge will be to exploit these differences to engineer SMC complexes that can topologically load onto DNA but not loop extrude, and *vice versa*. This will eventually enable genetic experiments that clarify the respective physiological contributions of topological loading and loop extrusion by SMC complexes to genome function.

## Material and methods

### Molecular model of the cohesin complex

The molecular model of cohesin in this study is based on our cryo-EM structure of the fission yeast cohesin complex together with its loader in the nucleotide-bound DNA gripping state (PDB: 6YUF) (*Higashi et al., 2020*). A molecular model of the fission yeast SMC hinge domain was obtained based on a mouse cohesin hinge crystal structure as a model (PDB:2WD5) (*Kurze et al., 2011*) using SWISS-MODEL (*Waterhouse et al., 2018*). Scc3^Psc3^ with the kleisin middle region, bound to DNA, was modeled based on a crystal structure of the corresponding budding yeast components (PDB: 6H8Q) (*Li et al., 2018*). The hinge and Scc3^Psc3^ were manually placed so that they align with the respective positions of the SMC hinge and Scc3^SA1^ in the cryo-EM structure of human cohesin in the gripping state (PDB: 6WG3) (*Shi et al., 2020*). The indicative position of Scc3^Psc3^ in the fission yeast structure, shown for comparison, was estimated based on distance constraints from a protein crosslink mass spectrometry dataset and guided by the negative staining EM density (*Higashi et al., 2020*). The coiled coils emanating from the ATPase heads were extended towards the SMC hinge using modeled Smc1^Psm1^ and Smc3^Psm3^ coiled coil segments, built based on their amino acid sequence using CCbuilder2.0 (*Wood and Woolfson, 2018*). The elbow positions in Smc1^Psm1^ and Smc3^Psm3^ were previously identified (*Higashi et al., 2020*).

To build a molecular model of cohesin in the post-hydrolysis slipping state, Scc2^Mis4^ and the Smc3^Psm3^ head domain were replaced with models of the same elements in new conformational forms, corresponding to previously observed free crystal structure states, as described (*Gligoris et al., 2014; Kikuchi et al., 2016; Higashi et al., 2020*). To model Brownian motion of the Scc3-hinge module relative to the Scc2-head module, Scc3^Psc3^ with the kleisin middle region and bound DNA together with the SMC hinge were considered to be one rigid body. The position of this Scc3-hinge module was then developed using a swinging motion of the SMC coiled coils with the inflection point at their elbows.

The molecular model of a nucleosome is based on the crystal structure of a human nucleosome (PDB: 3AFA) (*Tachiwana et al., 2010*). All structural figures were prepared using Pymol (Schrödinger) and ChimeraX (*Goddard et al., 2018*).

### Protein purification and labeling

For the construction of cohesin complexes harboring FRET reporters, SNAP- and CLIP-tag sequences were introduced into the YIplac211-Psm1-Psm3 and YIplac128-Rad21-Psc3 budding yeast expression vectors, as well as the pMis4(N191)-PA fission yeast expression vector that were previously described (*Murayama and Uhlmann, 2014; Chao et al., 2015; Higashi et al., 2020*). For labeling the Psm1 hinge, the SNAP tag sequence was inserted between Psm1 amino acids R593 and P594. Psc3 was fused to the SNAP or CLIP tag sequence at its C-terminus. For labeling Mis4, the CLIP tag sequence preceded the Mis4(N191) N-terminus. All proteins were purified and labeled with BG-surface Alexa 647 and BC-surface Dy547 dyes (New England Biolabs) as previously described (*Murayama and Uhlmann, 2014; Higashi et al., 2020*). The absorbance spectra of the labeled proteins were recorded between 220-800 nm in 1 nm increments using a V-550 Spectrophotometer (Jasco). The concentrations of protein, Dy547 and Alexa 647 were determined from the absorbance at 280, 550 and 650 nm, respectively, and the labeling efficiencies calculated (assuming molar extinction coefficient (ε) for Dy547 and Alexa 647 of 150,000 M^-1^ cm^-1^ and 270,000 M^-1^ cm^-1^, respectively).

### Bulk FRET measurements

Bulk FRET measurement were performed as previously described (*Higashi et al., 2020*), with minor modifications. All fluorescence measurements were carried out in reaction buffer (35 mM Tris-HCl pH 7.5, 0.5 mM TCEP, 25 mM NaCl, 1 mM MgCl2, 15% (w/v) glycerol and 0.003% (w/v) Tween 20). 40 μl of reaction mixtures containing 37.5 nM of the respective labeled cohesin, 100 nM Mis4-Ssl3 or 12.5 nM Dy547-labeled Mis4(N191) and 10 nM pBluescript KSII(+) as the DNA substrate were mixed and the reaction was started by addition of 0.5 mM ATP. Alternatively, 0.5 mM ADP and 0.5 mM BeSO4 + 10 mM NaF were included to generate the DNA gripping state. The reactions were incubated at 32 °C for 20 minutes. The samples were then applied to a 384-well plate and fluorescence spectra were recorded at 25 °C on a CLARIOstar Microplate Reader (BMG LABTECH). Samples were excited at 525 nm and emitted light was recorded between 560 - 700 nm in 0.5 nm increments. To evaluate FRET changes caused by cohesin’s conformational changes across different experimental conditions, we report relative FRET efficiency, IA/(ID + IA), where ID is the donor emission signal intensity at 565 nm resulting from donor excitation at 525 nm and IA is the acceptor emission signal intensity at 665 nm resulting from donor excitation. To obtain a baseline of apparent FRET due to spectral overlap, we mixed singly Dy547 and Alexa 647-labeled cohesin at concentrations of 12.5 nM and 37.5 nM, respectively, to reflect the fluorophore stoichiometry of the double labeled complex.

### Co-immunoprecipitation

100 nM cohesin labeled at the Psm1 hinge with Alexa 647, 100 nM Mis4ΔN191 labeled at the N-terminus with Dy547, 10 nM pBluescript KSII (+) DNA and 0.5 mM ATP or 0.5 mM ADP/0.5 mM BeSO4/10 mM NaF were incubated in 15 μl of reaction buffer at 32 °C for 20 minutes then diluted with 100 μl of reaction buffer. 5 μl of anti-Pk antibody (Bio-Rad, MCA1360)-bound Protein A conjugated magnetic beads were added to the reactions and rotated at 25 °C for 10 minutes. The beads were washed twice with 500 μl of reaction buffer and the recovered proteins were analyzed by SDS-PAGE followed by in gel fluorescence detection, as well as immunoblotting with the indicated antibodies.

### Mathematical modeling of loop extrusion

#### DNA model

We based modeling of our DNA on a dssWLC model, as described in (*Koslover and Spakowitz, 2014*). The DNA is defined by a sequence of beads with positions 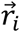and an orientation unit vector 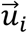 attached to each bead. The energy for a particular chain configuration is given by:

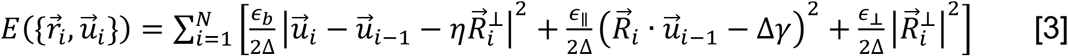

Where 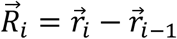 and 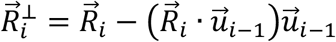

The DNA in this model is split into segments with contour length Δ. The model parameters *∈*_*b*_, *∈*_*∥*_, *∈*_⊥_, *γ* and *η* are unambiguous functions of Δ and polymer persistence length *l*_*p*_ taken from (*Koslover and Spakowitz, 2013*). For all our simulations we use Δ = 5 nm and *l*_*p*_ = 50 nm.

#### Cohesin model

We generated a simplified model of cohesin using five beads representing the Smc1 and Smc3 heads, the Smc1 and Smc3 elbows as well as the hinge. Similar to DNA, each bead is defined by its position and orientation vectors (e.g. 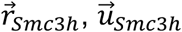 for the Smc3 head, *etc*.). We describe the interaction between hinge, elbow and the corresponding head beads via dssWLCs and the corresponding energy term is given by [3]. Based on the experimentally determined elbow position (*Bürmann et al., 2019; Higashi et al., 2020*), we choose contour length between the head and elbow to be 30 nm, while the contour length between the elbow and hinge is 20 nm. In order to determine the persistence length of the connecting coiled coil for our simulations, we sampled conformational states of cohesin alone as it transitions multiple times between slipping and gripping states. We found that at the persistence length *l*_*cc*_ = 50 nm our simulations match the experimentally observed head-to-hinge distance distribution available for the condensin complex (*Ryu et al., 2020b*). A smaller persistence length for condensin’s coiled coil of *l*_*cc*_ = 4 nm was previously reported (*Eeftens et al., 2016*), which included the flexible elbow region that is considered separately in our simulations.

The interaction between the Smc1 and Smc3 heads is treated differently because of their geometry. We describe their interaction with the same dssWLC approach, corrected for their positions and orientations. Interaction energy between the head domains is given by:

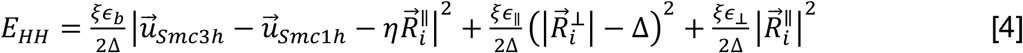

Where 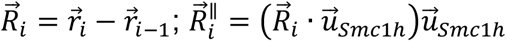 and 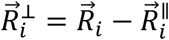.

In [4] we use the same set of parameters *∈*_*b*_, *∈*_*∥*_, *∈*_⊥_, *γ* and *η* as in [3] for the SMC coiled coils. However, SMC heads are connected by the cohesin loader subunit, which provides unknown additional stiffness to this connection. To take this into account, we introduce an additional parameter *ξ*. We find that as long as *ξ* > 5, it does not noticeably affect our simulations. Therefore, for most simulations we used a value of *ξ* = 10.

#### Interaction between Cohesin and DNA

We introduce two separate energy terms describing interactions between DNA and the Smc3 head as well as DNA and the hinge.

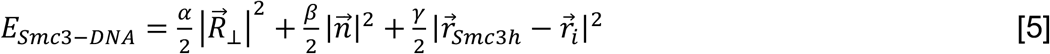

Where 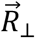 is the shortest distance between Smc3 and the center of the closest DNA bead,

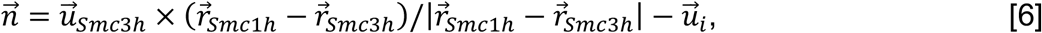

where *i* is the index of the DNA bead currently interacting with Smc3.

In [5], the first two terms describe the slipping interaction between DNA and Smc3 which allows diffusion of DNA along the Smc3 head. The second term is introduced to take into account the orientation of DNA with respect to the orientation of cohesin. The third term describes a point-to-point gripping interaction. We assume *γ* = 0 in the slipping state.

The interaction between the DNA and the hinge is:

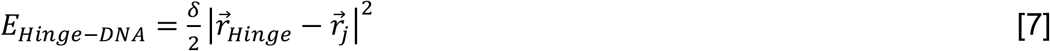

Where *j* is the index of the DNA bead currently interacting with the hinge.

[5] and [7] describe physical bonds between DNA and cohesin. Because these terms grow indefinitely with the relative distance between the two, they do not describe the dynamics of the bond breakage. In order to account for cohesin changes between gripping and slipping states, parameter *γ* was changed to zero for the slipping state. The exact values of parameters *α, β, γ* and *δ* are inconsequential as they only affect the extent to which the relative distance between cohesin and the DNA can fluctuate. We have chosen values of these parameters as a trade-off between minimizing fluctuations to within one spatial discretization step (5 nm) and improving algorithm convergence.

#### 3D Monte-Carlo simulations

Numerical calculations were carried out with the Metropolis method for Monte-Carlo simulation (*Heermann, 1990*). Briefly, random beads representing a DNA segment or a part of cohesin is chosen at each step and its position or orientation vector is randomly modified. The new full energy of the system *E*_*new*_ is calculated. If the new energy is lower than the previous value *E*_*new*_, the new state is accepted. If it is larger, the new state Boltzmann factor is calculated: 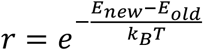 and the new state is accepted if *r* > *p*, where *p* is drawn from a standard uniform distribution on the interval (0,1).

#### Simplified model of diffusion-based loop extrusion

In the simplified model for DNA loop extrusion, a DNA loop inside a cohesin molecule consists of N segments, each 5 nm in length. On each iteration of the algorithm, we consider all possible events that can happen to the system consisting of cohesin and a DNA loop inside. There are five types of events: 1. Random thermal movement of the inbound DNA at the Smc3 head (only possible in the slipping state), 2. Random thermal movement of the outbound DNA, which does not interact with Smc3, 3. If the hinge is bound to DNA, DNA can move with the hinge if cohesin changes its state from the folded gripping to the unfolded slipping state, or the other way around, 4. The hinge can bind/unbind DNA, 5. Cohesin can change state, from gripping to slipping state, or *vice versa*.

At each iteration, rate constants are calculated for all possible events. The rate constant of DNA diffusion is the rate at which one DNA segment moves one step forward or backward. It is given by:

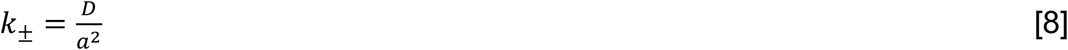

Where *D* is the diffusion coefficient and *a* = 5 *nm* length of the DNA segment.

Off-rate of hinge DNA unbinding:

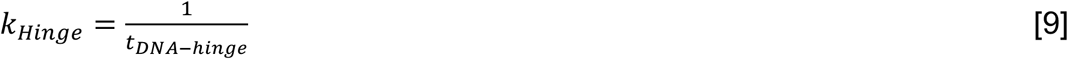

Where *t*_*DNA-hinge*_ is the lifetime of DNA-hinge interaction defined in the main text.

We assume that the rates of slipping -> gripping and gripping -> slipping state conversion are the same:

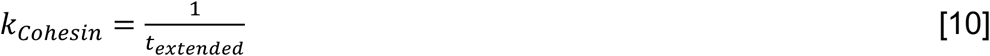

Where *t*_*extended*_ is the lifetime of the slipping state defined in the main text.[10]

At a given iteration of the Monte Carlo simulation, we calculate a set of times at which each possible event stochastically occurs:

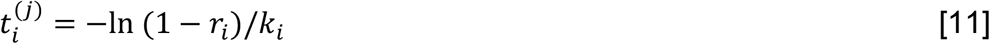

Where *j* is the current iteration, *i* is the given event. *r*_*i*_ is a uniformly distributed number on the interval (0,1), and *K*_*i*_ the rate for event *i*. Then we implement the event which has the smallest 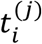 and modify the system based on which event occurred. The total time of the simulation is extended by 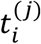.

#### Estimation of the diffusion-driven rate of cohesin unfolding

The simplest assumption about the nature of the cohesin transition from the folded gripping to the unfolded slipping state is that it is driven by diffusion only. The diffusion coefficient can be estimated as:

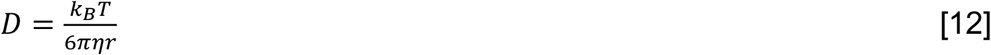

where *η* = 10^−3^ Pa·s is the viscosity of water and *r* is the effective radius of the diffusing protein. The estimate of the time it takes to diffuse distance *x* is:

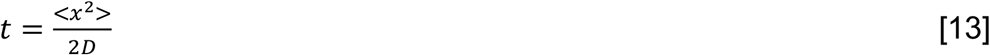

The size of the protein is related to its molecular weight. Given cohesin’s irregular shape, the diffusion can be better estimated based on the actual size of the protein. Using a size of ∼ 20 nm for the Scc3-hinge and Scc2-head modules, we get a conservative estimate of ∼ 0.1 ms for the time it will take the modules to separate by diffusion ∼ 50 nm.

## Acknowledgements

We thank A. Costa and S. Henrikus for their collaboration on cohesin structural biology and for their advice, S. Kunzelmann and S. Mouilleron from the Crick Structural Biology Science Technology Platform for instrument access and advice and all our laboratory members for discussions and critical reading of the manuscript.

## Additional information

### Funding

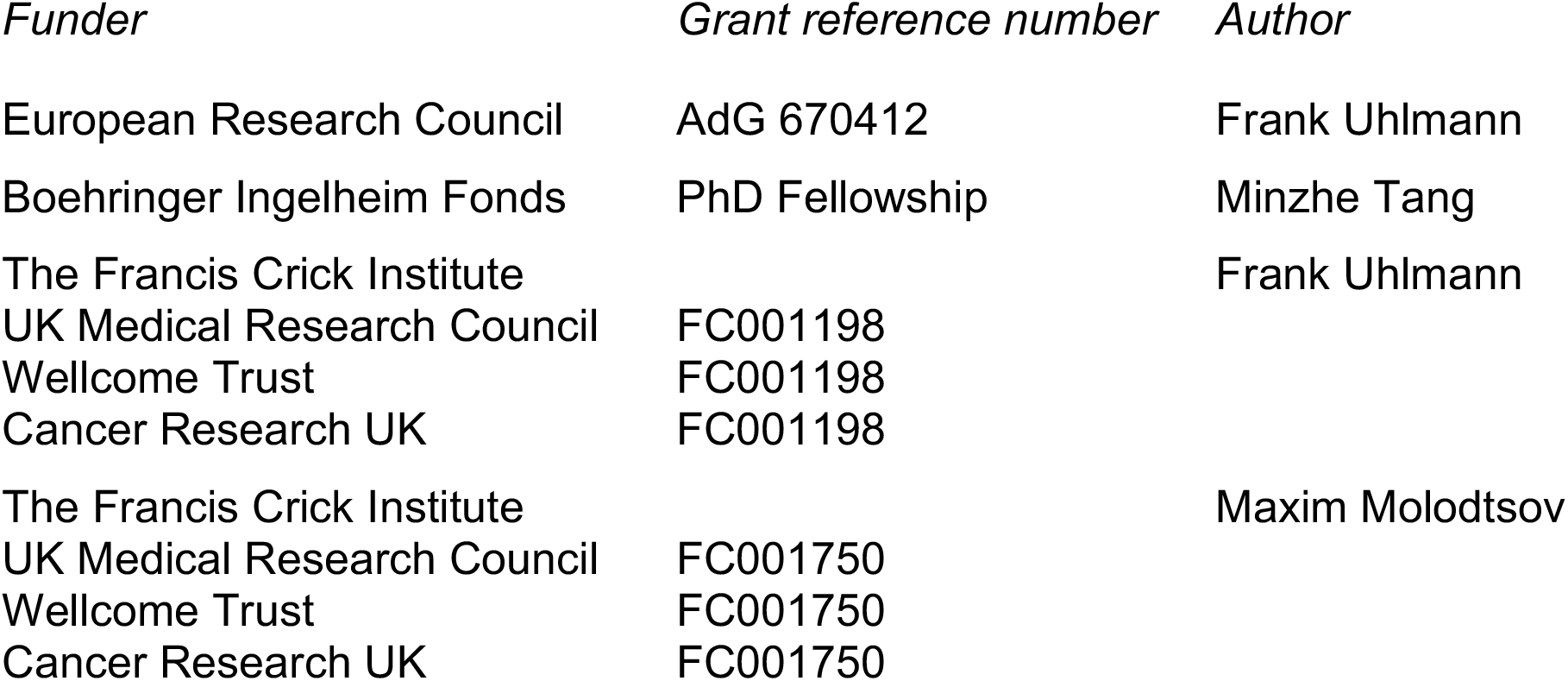

The funders had no role in study design, data collection and interpretation, or the decision to submit the work for publication.

### Author Contributions

T.L.H., F.U. and M.M. conceived the study, T.L.H. performed the experiments and drew the structural figures, M.T. contributed the concept of second DNA capture by condensin, G.P. and M.M. designed the mathematical model and performed the computational simulations. T.L.H., F.U. and M.M. wrote the manuscript with input from all coauthors.

---

**Figure 1 - figure supplement 1.**
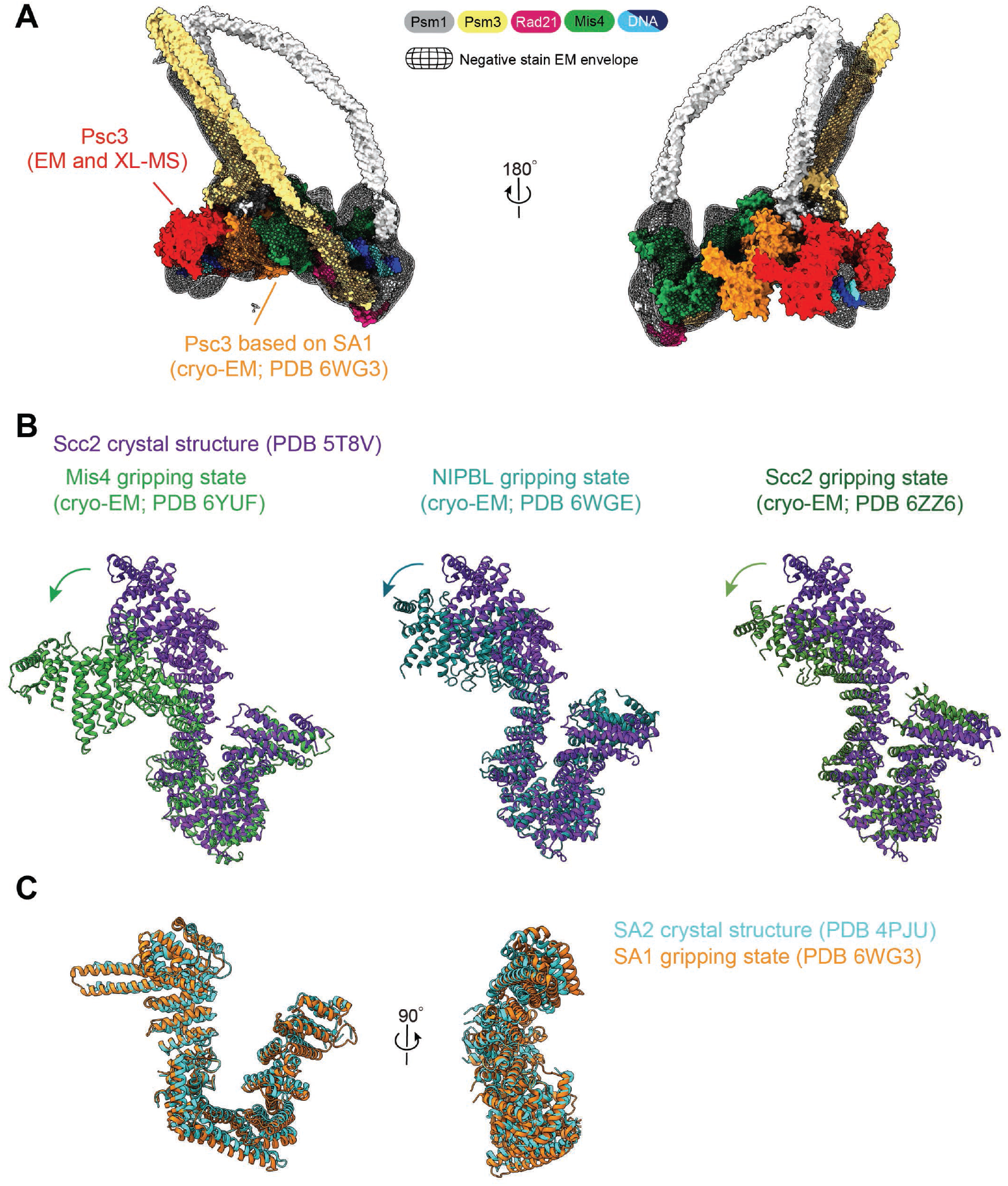
Configuration and conformations of Scc3^Psc3^ and Scc2^Mis4^ in the DNA gripping state. **(A)** Molecular overview model of the DNA gripping state, highlighting the Scc3^Psc3^ configuration. The position of Psc3 based on our negative stain EM and protein crosslink mass spectrometry (XL-MS) analyses is shown in red. The configuration based on the position of its SA1 paralog in the cryo-EM structure of human cohesin (*Shi et al., 2020*) is shown in orange. **(B)** Structural comparisons of Scc2^Mis4^ in the fission yeast, human and budding yeast gripping states (*Collier et at., 2020; Higashi et al., 2020; Shi et al., 2020*), aligned to the extended C. *thermophilum* Scc2 crystal structure form (*Kikuchi et al., 2016*). **(C)** Structural comparison of human Scc3^SA1^ in the gripping state with the free crystal structure form of its paralog Scc3^SA2^ (*Hara et al., 2014*).

**Figure 2 - figure supplement 1.**
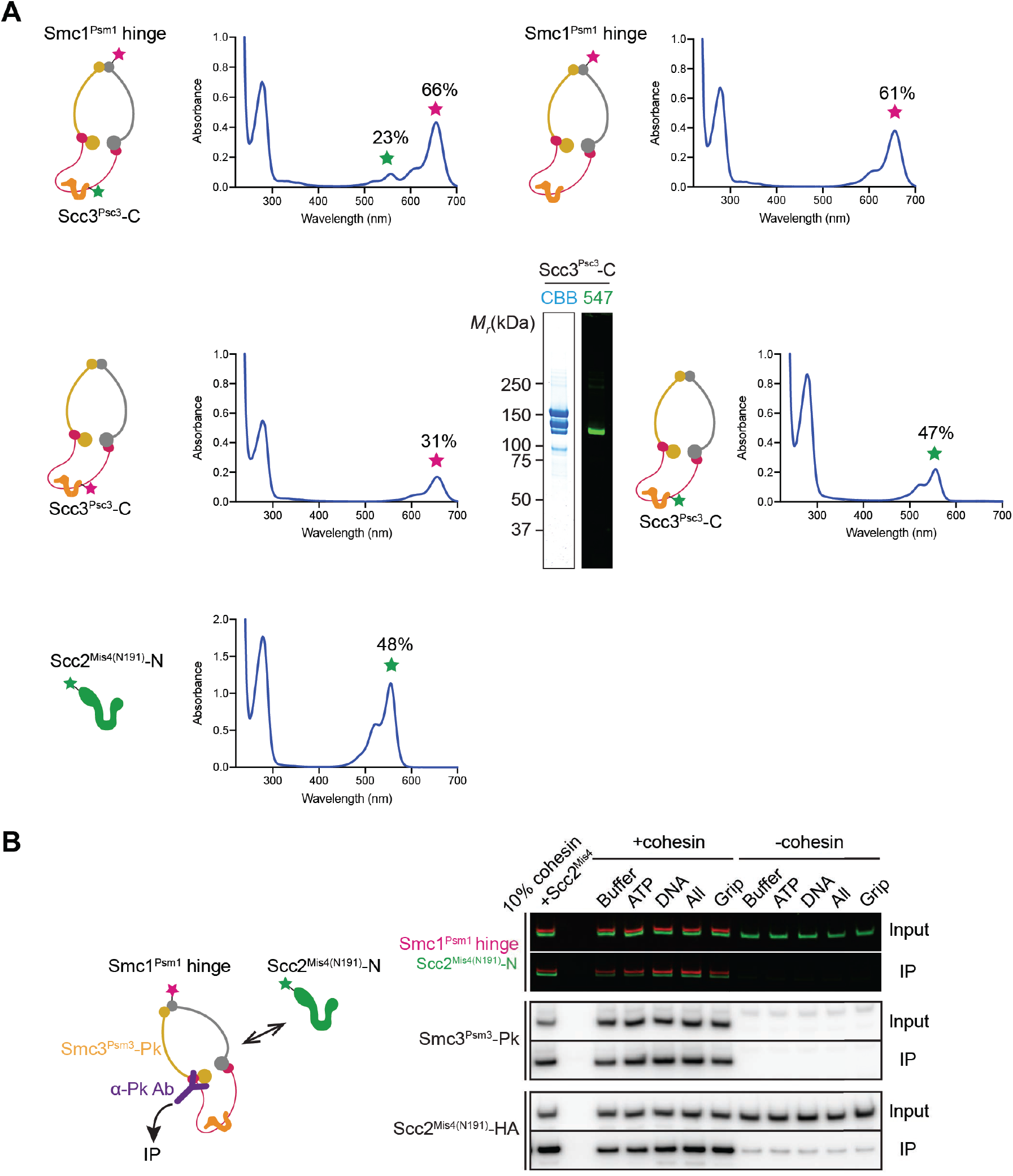
Supporting control experiments for the FRET-based conformational analyses. **(A)** Labeling efficiencies of the cohesin complexes and Scc2^Mis4(N191)^, calculated based on protein absorbance at 280 nm and fluorophore absorbance at 560 and 660 nm as detailed in the Material and methods. Shown is also the purified cohesin complex, single-labeled with Dy547 at the Scc3^Psc3^ C-terminus, used for the FRET baseline determination, analyzed by SDS-PAGE followed by Coomassie blue staining (CBB) and in gel fluorescence detection. Labeling efficiencies were typically around 50%, though somewhat lower in the case of Psc3, maybe due to the tendency of the latter subunit to be a substoichiometric component of the cohesin complex during protein purification. **(B)** Investigation of complex formation between cohesin and the Scc2^Mis4(N191)^ cohesin loader. Following incubation of cohesin and Scc2^Mis4(N191)^ under the conditions used for FRET recordings, cohesin was immunoprecipitated via a Pk epitope tag at the Smc3^Psm3^ C-terminus. Coprecipitation of Scc2^Mis4(N191)^ was analyzed by SDS-PAGE followed by immunoblotting and in-gel fluorescence detection.

**Figure 3 - figure supplement 1.**
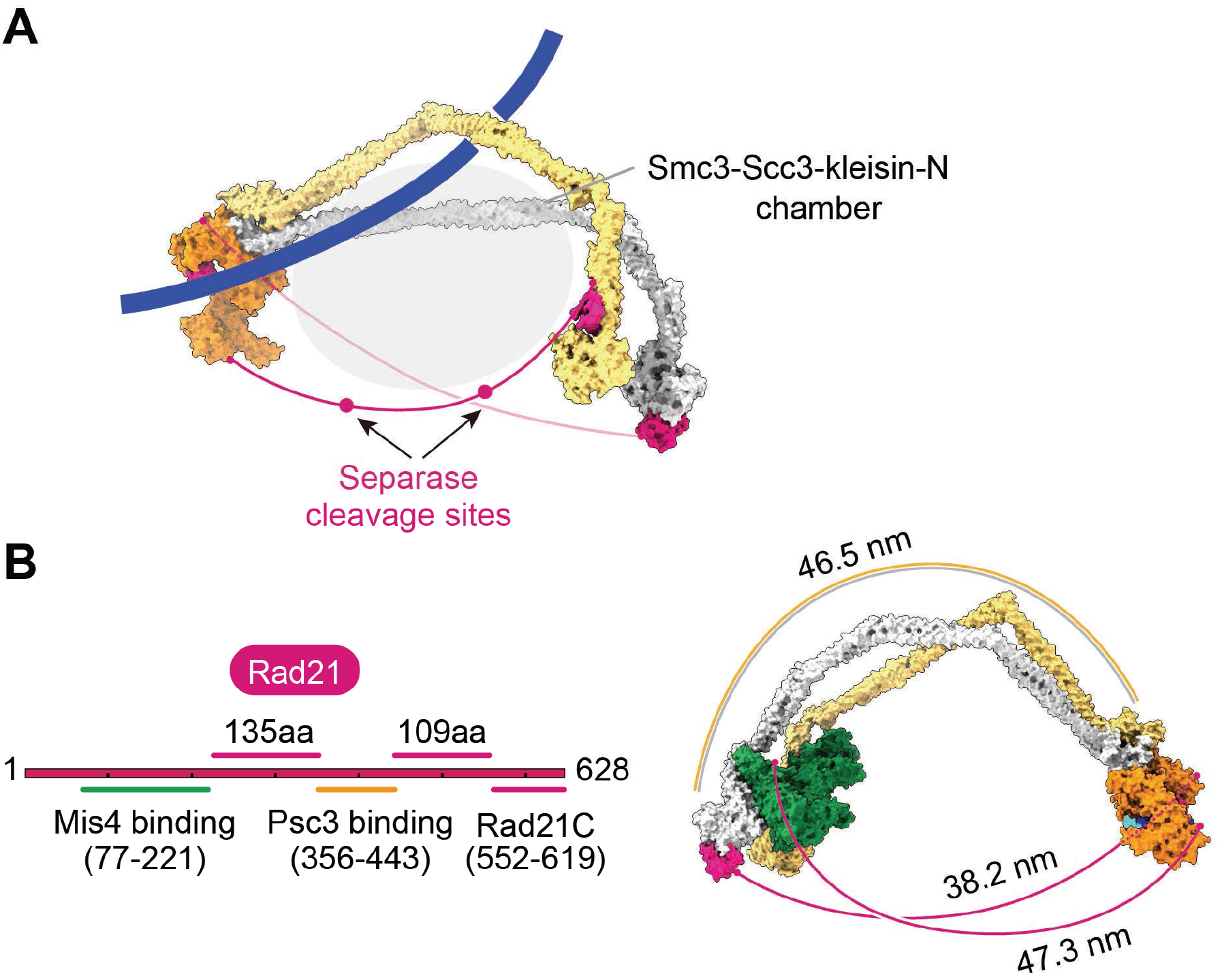
Additional views of cohesin during topological loading onto DNA and during loop extrusion. **(A)** Following completion of topological cohesin loading, DNA finds itself in the Smc3-Scc3-kleisin-N chamber, delineated by the Smc3^Psm3^ coiled coil, Scc3^Psc3^ and the unstructured kleisin region between its Scc3^Psc3^ interacting sequences and the kleisin N-gate. Two separase recognition sites, whose cleavage liberates cohesin from DNA to trigger anaphase, are indicated. **(B)** Schematic of the kleisin and the lengths of its unstructured regions. A molecular model depicts the potential span of the kleisin, using a conservative estimate of 3.5 Å per amino acid (*Ainavarapu et al., 2007*), compared to the length of the SMC coiled coil, after the Scc3-hinge module separates from the Scc2-head module during loop extrusion.

**Figure 5 - figure supplement 1.**
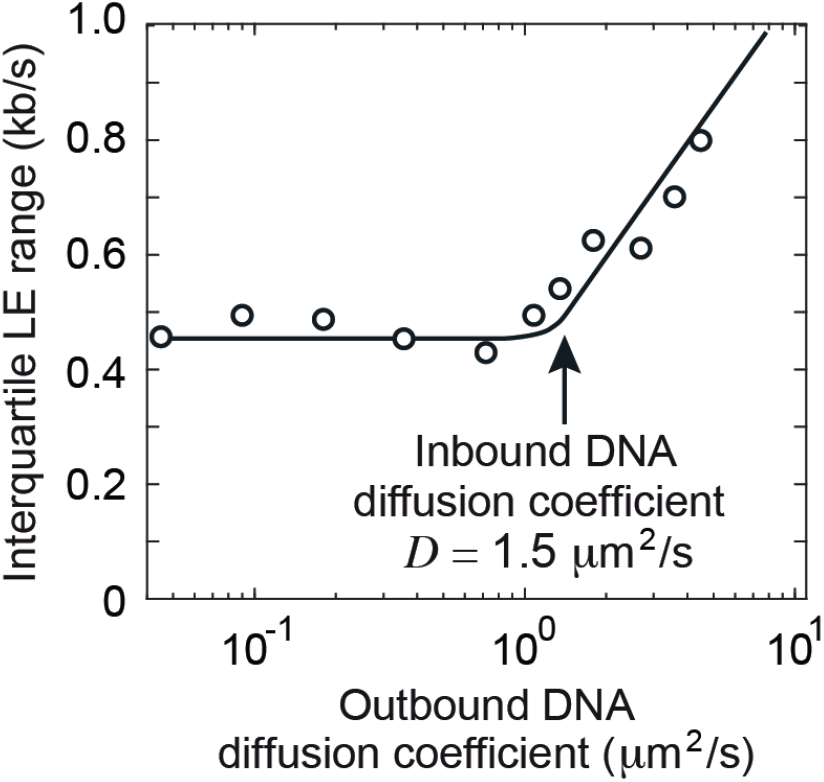
Interquartile range of loop extrusion (LE) rates when the DNAs entering and exiting the cohesin ring show divergent diffusion coefficients. The diffusion coefficient of the inbound DNA was fixed at 1.5 μm^2^/s, while the diffusion coefficient of the outbound DNA varied between 0.05 and 5 μm^2^/s. The line indicates the overall trend. Simulations were over 3 minutes.

**Figure 7 - figure supplement 1.**
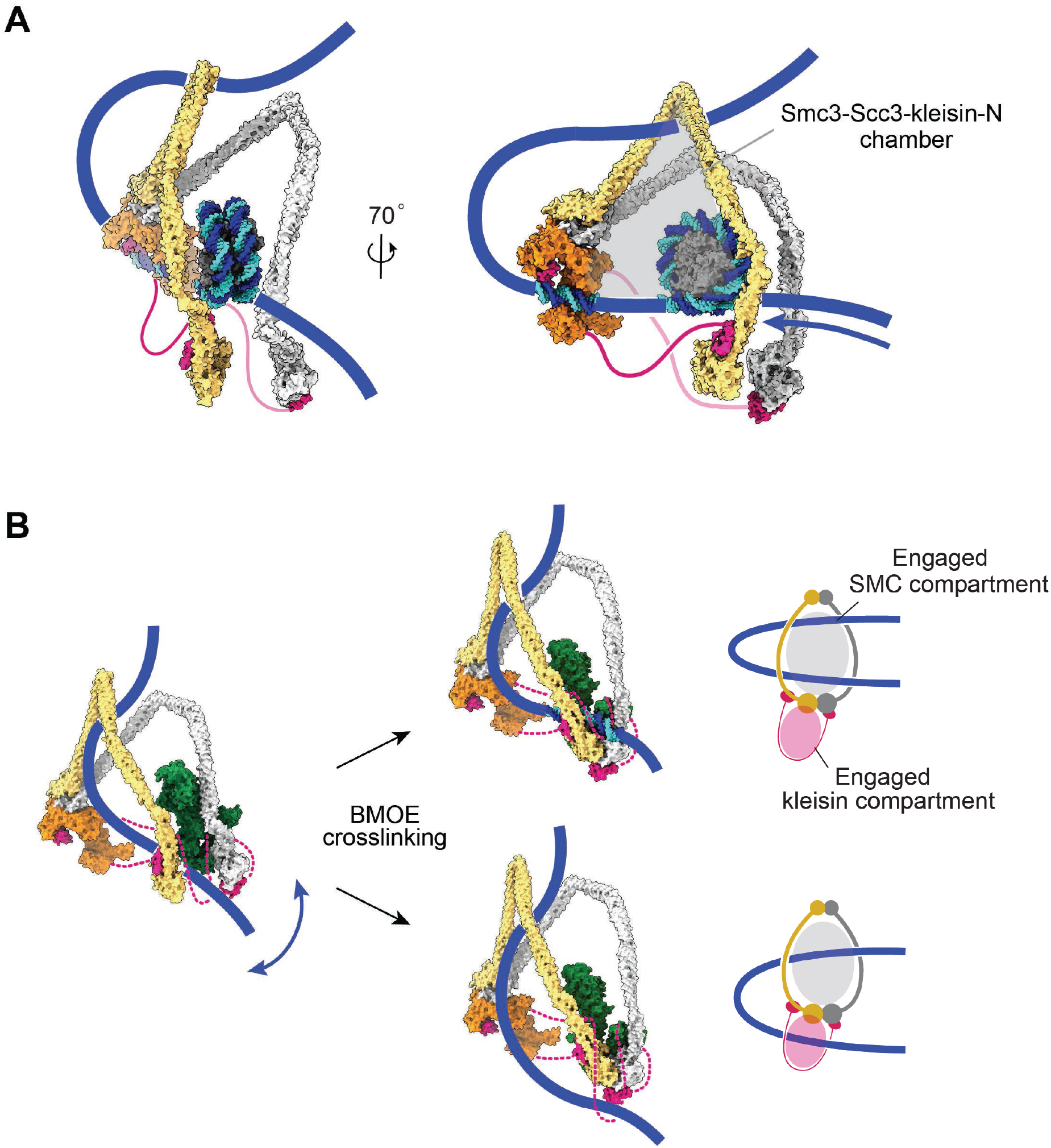
Additional views of cohesin during nucleosome bypass and during the early stages of encountering DNA. (**A**) Molecular model of nucleosome bypass by cohesin. Following dissociation of the cohesin loader, directed Brownian motion pulls incoming DNA through the Smc3-Scc3-kleisin-N chamber. This chamber is wide enough to accommodate a nucleosome and possibly even larger DNA-bound obstacles. (**B**) Schematic that reconciles our molecular model of cohesin function with observations of the DNA position within the cohesin ring based on chemical crosslinking (*Collier et al., 2020*). Our DNA-protein crosslink mass spectrometry data suggest that DNA initially explores the space between the SMC coiled coils. Before ATP-dependent head engagement, this might include passage between the heads. BMOE-induced crosslinking results in two possible outcomes. DNA is either not trapped by cohesin (top), or DNA is trapped in both the engaged SMC compartment, as well as the engaged kleisin compartment (bottom). At a slower rate, DNA successfully passes the kleisin N-gate upon ATPase head engagement, leading to topological DNA entry into the SMC-kleisin ring.

**Supplementary video 1**. Progression of the Metropolis Monte-Carlo simulation as cohesin switches between gripping and slipping states. Before the start of the video, the system was equilibrated for ∼ 10^7^ iterations. Frames were captured every 2×10^5^ iterations and the video spans ∼5×10^7^ iterations. Pauses were introduced to highlight state transitions and DNA binding and unbinding events. Axis scale, nm.

